# Enhancing PDAC Therapy: Decitabine-Olaparib Synergy Targets KRAS-Dependent Tumors

**DOI:** 10.1101/2024.06.23.599676

**Authors:** Giorgia Anastasio, Michela Felaco, Alessia Lamolinara, Francesco del Pizzo, Elisa Cacciagrano, Carla Mottini, Margherita Mutarelli, Francesca Di Modugno, Manuela Iezzi, Luca Cardone

**Affiliations:** Institute of Biochemistry and Cellular Biology, National Research Council (CNR), Rome, Italy; Unit of Tumor Immunology and Immunotherapy, IRCCS Regina Elena National Cancer Institute, Rome, Italy; Center for Advanced Studies and Technology (CAST), Chieti, Italy; Department of Neurosciences, Imaging and Clinical Sciences, G. d’Annunzio University of Chieti-Pescara, Chieti, Italy; UOSD SAFU Translational Research Area, IRCCS Regina Elena National Cancer Institute, Rome, Italy; Institute of Applied Sciences and Intelligent Systems (ISASI), CNR, Naples, Italy

## Abstract

Current chemotherapies provide limited clinical benefits to patients with pancreatic ductal adenocarcinoma (PDAC), partly due to the lack of effective biomarkers for personalized therapy. KRAS activating mutations occur in almost 90% of PDAC cases, leading to a subset of tumors <u>d</u>ependent on <u>KRAS</u> for survival (dKRAS). Assessing dKRAS in PDAC can be achieved using gene expression signature scores, independent of specific KRAS mutations, allowing for personalized therapies. Previous studies have shown that dKRAS-PDAC cells are more sensitive to the FDA-approved drug decitabine (DEC), although the mechanism remains unclear. While DEC is approved for hematological tumors, its repurposing in solid tumors poses challenges due to high-dose hematological side effects. Identifying optimal pharmacological approaches and response biomarkers is crucial for the successful clinical implementation of DEC in PDAC and other solid tumors. Our investigation revealed that low-dose DEC combined with the PARP inhibitor olaparib (OLA) enhances antitumor activity in dKRAS-PDAC. Mechanistically, DEC induces DNA damage and activates an ATR/ATM-mediated DNA damage response (DDR), with PARP1-mediated DNA repair playing a crucial role. Inhibiting PARP activity with OLA enhances antitumor activity, even in *BRCA1/2*^*wild-type*^ and homologous recombination (HR)-proficient tumors, but it is ineffective in KRAS-independent tumors. Thus, transcriptomic-based KRAS dependency scores effectively predict the efficacy of combined therapy. Additionally, in dKRAS-PDAC tumors carrying a *BRCA2* mutation, low-dose DEC enhances OLA’s antitumor activity, completely inhibiting metastasis growth compared to single-drug treatments. Our findings support further clinical evaluation of DEC+OLA combination therapy in PDAC, particularly in dKRAS-positive tumors, irrespective of BRCA1/2 status. This approach extends the clinical benefit of OLA beyond BRCA status, addressing a limitation in targeted PARP inhibitor therapies. Furthermore, we highlight DDR as a key mechanism of action for DEC, beyond its role in gene expression regulation, underscoring the mode of action of this widely used anticancer drug.

## INTRODUCTION

With a 5-year survival rate of only 8%, pancreatic ductal adenocarcinoma (PDAC) is the fourth leading cause of cancer death worldwide ^1^. Despite extensive efforts, there has been limited improvement in the therapeutic and clinical management of advanced and metastatic PDAC, as chemoresistance and tumor recurrence rates remain high. Reliable diagnostic markers for personalized therapy are still unavailable, resulting in treatments being administered without stratifying patients for optimal benefits and likelihood of response.

Between 90–95% of PDAC cases harbor a KRAS oncogene-activating mutation ^2^. Genetically modified mouse models have demonstrated the essential role of KRAS-dependent signaling in PDAC growth and development ^3^. By investigating KRAS-associated gene profiles in PDAC, subtypes with a molecular <u>d</u>ependency on <u>KRAS</u> (dKRAS) can be identified, referring to tumors whose growth still requires active KRAS-dependent signaling ^4,5^. The extent of KRAS dependency can differ even with a similar genomic KRAS mutation ^4,5^, indicating that KRAS mutations might have limited predictive value regarding tumor cells’ reliance on KRAS-dependent pathways. Computational gene signature-based scores can stratify tumors, cancer cell lines, and patient-derived xenografts (PDX) to predict tumor cell dependence on KRAS (dKRAS) in PDAC. A significant proportion of PDAC cases fall under the dKRAS subtype ^4^. Consequently, transcriptional scores of KRAS activation and dependency can be effectively exploited as biomarkers for tumor classification in PDAC.

Drug repurposing, the use of FDA-approved medicines for new therapeutic indications, is a novel approach to redirecting therapies in cancer ^6,7^. Decitabine (5-Aza-2-deoxycytidine, DEC) is approved to treat myelodysplastic syndromes and acute myeloid leukemia. Its anticancer activity has been considered in other clinical settings, with more than fifty clinical trials testing the repurposing of DEC alone or in combination with chemotherapies or immunotherapies for treating solid tumors, including breast, ovarian, colon, lung, pancreatic cancers, and sarcoma ^8,9^. However, DEC treatment is associated with severe hematological toxicity in approximately 30% of cases, including G3/4 febrile neutropenia. These adverse events may limit treatment efficacy and adversely affect the quality of life for patients, particularly with high-dose administration ^10^. This underscores the importance of identifying therapeutic strategies to mitigate DEC side effects while preserving its anticancer efficacy. Enhancing our understanding of DEC’s mode of action and identifying biomarkers that predict patient response to DEC is crucial for its successful clinical repurposing in solid tumors.

DEC, an irreversible inhibitor of DNA methyltransferase and a cytosine analog incorporated into DNA during replication, has been extensively studied for its anticancer activity as a DNA-demethylating agent ^11^. This activity results in gene expression modulation, specifically reactivation of antiapoptotic genes ^12^ enhancing immune-checkpoint inhibitor therapy by overexpressing mutated neoantigens ^13^, or reverting T-cell exhaustion ^14^. We previously demonstrated that DEC exhibited potent antitumor and antimetastatic properties in the KRAS-dependent PDAC subtype, although the mechanism remains unclear. In dKRAS-but not in KRAS-independent PDAC, DEC treatment induced cell-cycle arrest at the G2/M phase and induced phospho-H2AX, a DNA damage response (DDR) marker ^7^, suggesting that, in dKRAS-PDAC, DEC might induce DNA damage. Research findings have shown that DEC can induce a DNA damage response mediated by the Ataxia Telangiectasia Mutated (ATM) protein and lead to double-strand breaks in HeLa cells, colon cancer cells ^15^, and multiple myeloma cells ^16^. Moreover, acute myeloid leukemia cells deficient in base excision repair (BER) were hypersensitive to DEC, and BER was required to remove aberrant bases from DNA and recognize abasic sites generated by DEC ^17^. Thus, the available evidence suggests that DEC’s mechanism of action as an antitumor agent might also involve DDR.

The Poly (ADP-ribose) polymerase (PARP) plays a key role in DNA damage response (DDR) to maintain genomic integrity in highly replicative cells. PARP inhibitors (PARPi) have demonstrated clinical efficacy against several solid tumor types carrying BRCA1/2 genomic mutations and homologous recombination deficiency (HRD), through a synthetic lethal-based mechanism ^18,19^. In PDAC, the PARPi olaparib (OLA) has been approved as maintenance therapy in platinum-treated tumors carrying BRCA1/2 mutations. More than forty clinical trials are testing the efficacy of PARPi alone or in combination with other treatments at various stages of PDAC. There is also growing interest in investigating the clinical efficacy of PARPi in solid tumors regardless of BRCA1/2 mutations, based on functional deficits in BRCA (BRCAness), microsatellite instability (MSI), or defects in the DNA homologous recombination repair (HRR) pathway. Genomic mutational profiles of HR-related genes are under investigation as biomarkers of response ^20^. Additionally, molecular surrogate readouts of HRD can identify a greater proportion of patients with HRD than analyses limited to gene-level approaches ^20,21^. Evidence in breast and ovarian cancers has demonstrated that a RAD51 score, estimated as the percentage of geminin-positive cells carrying RAD51-positive nuclear foci, might predict tumor cellular HR status. RAD51 scores are very low in HR-deficient cells and correlate with responses to PARPi or platinum treatments ^22^. Thus, extending the population who might benefit from PARPi beyond the available biomarkers, which is also the focus of the present manuscript, could significantly enhance clinical practice and treatment options for PDAC patients.

Previous studies have demonstrated molecular crosstalk between PARP activity and DEC. OLA has been demonstrated to block DEC-induced BER replication fork collapse, thereby inducing double-strand breaks and cell death, and promoting synthetic lethality in acute myeloid leukemia (AML) ^17^. Further research has reported increased cytotoxic effects of combining PARP inhibitors with DNA demethylating agents in AML subtypes and BRCA wild-type breast cancer cells ^23^, particularly in combination with irradiation. Based on this evidence, we speculated that PARP activity is involved in the repair process following DEC-induced DNA damage in KRAS-dependent PDAC (dKRAS-PDAC), potentially providing a therapeutic window for a synergistic drug combination of DEC with PARP inhibitors such as OLA. We investigated the preclinical efficacy of the combination of DEC and OLA in the dKRAS-PDAC subtype identified using transcriptional-based scores for KRAS dependency.

## RESULTS

### DEC induces robust DNA damage and DDR activation in KRAS-dependent PDAC cells through ATM/ATR-mediated pathways

PDAC cellular models from the CCLE panel can be classified based on transcriptomic assessments of their similarity to reference gene signatures of KRAS dependency and activation (referred to as the S- and L-scores, respectively), previously validated (Figure S1 and Table S1). To elucidate the mode of action of DEC in PDAC, we investigated the activation of DNA damage response (DDR) markers associated with single-strand breaks (SSBs) and double-strand breaks (DSBs). This included examining XRCC1-positive nuclear foci for SSBs and 53BP1- and RAD51-positive nuclear foci for DSBs. We selected dKRAS-PDAC cell lines such as BxPC-3, PaTu 8902, HPAF-II, and CAPAN-1 based on their high KRAS dependency scores (both L- and S-scores) (Figure S1 and Table S1). Quantification of immunofluorescence images showed a strong and statistically significant induction of phospho-Ser139(γ)H2AX-, XRCC1-, RAD51-, and 53BP1-positive nuclear foci in dKRAS-PDAC cells following 72 hours of DEC treatment. In contrast, the analysis of KRAS-independent (indepKRAS) cell lines, those with low transcriptional scores for KRAS dependency and activation (Figure S1 and Table S1), such as PaTu 8988t and KP-4, did not show any significant increase in number of nuclear foci for these markers following DEC treatment (Figure 1A, Figure S3). Furthermore, γH2AX-positive foci induced by DEC in dKRAS-PDAC cells colocalized with RAD51- and 53BP1-positive foci (Figure 1B).

**Figure 1.**
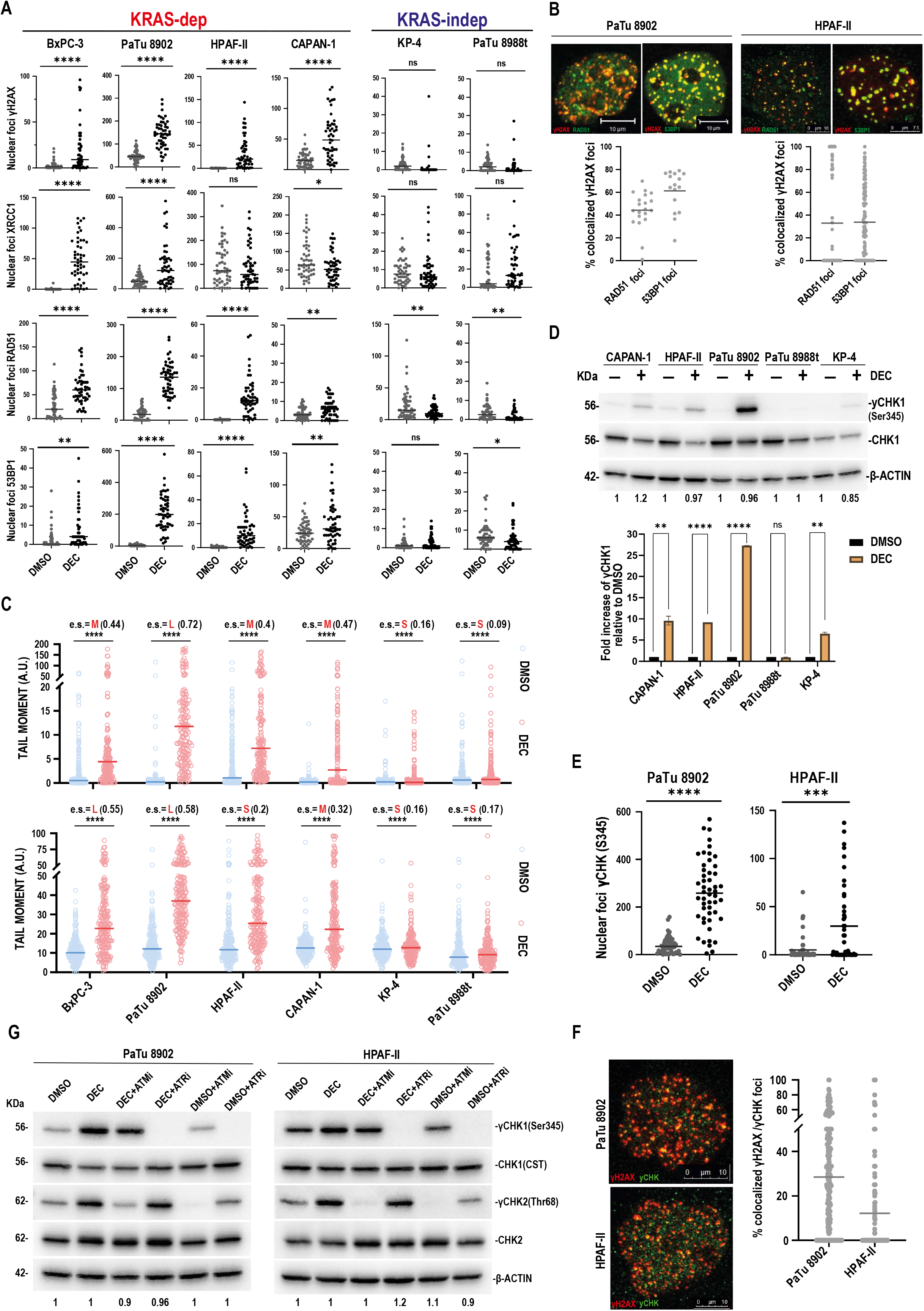
DEC induces robust DNA damage and DDR activation in KRAS-dependent PDAC cells through ATM/ATR-mediated pathways. (A) Quantification of immunofluorescence analysis of phospho-(γ)H2AX(Ser139)-, XRCC1-, RAD51-, and 53BP1-positive nuclear foci in dKRAS-PDAC cell lines (BxPC-3, PaTu 8902, HPAF-II, and CAPAN-1) and indepKRAS-PDAC cell lines (PaTu 8988t and KP-4) following DMSO (vehicle) or DEC (1.25 μM) treatment for 72 hours. Data are presented as median from a representative replicate experiment. At least 50 nuclei were counted for each replicate using immunofluorescence analysis with CellProfiler 4.2.1. Statistical analysis was performed using GraphPad Prism. Statistical significance for DMSO vs. DEC-treated cells was calculated using the unpaired two-tailed Student’s t-test: *p < 0.05, **p < 0.01, ***p < 0.001, ****p < 0.0001, ns: p > 0.05. For representative images of the experiment, please see Figure S3. (B) Colocalization analysis of γH2AX with RAD51- and 53BP1-positive nuclear foci in dKRAS-PDAC cells following DEC (1.25 μM) treatment for 72 hours. Upper panels: Representative images of confocal microscopy analysis of PaTu 8902 and HPAF-II co-stained for γH2AX with RAD51 or 53BP1. The nuclei were counterstained with DAPI. Scale bar, 10 μm. Magnification, x60. Lower panels: Dot plots representative of the percentage of colocalized foci. Data are presented as median from a representative replicate experiment. From 20 to 100 nuclei were counted for each replicate using immunofluorescence analysis with CellProfiler 4.2.1. Data analysis was performed using GraphPad Prism. (C) Quantification of comet tail moment (TM), which measures DNA fragmentation, in dKRAS-PDAC cells (PaTu 8902, BxPC-3, HPAF-II, and CAPAN-1) and indepKRAS-PDAC cells (PaTu 8988t and KP-4) following treatment with DMSO (vehicle) or DEC (1.25 μM) for 72 hours. Data are presented as median from a representative replicate experiment. At least 200 nuclei were counted for each replicate. Alkaline (upper panel) and neutral (lower panel) comet assays were performed. To quantify the magnitude of DEC-induced DNA damage compared to DMSO, the effect size (e.s.) was estimated by the Wilcoxon test from three to six biological replicates. The Rstatix package was implemented for the calculation of the effect size. Statistical significance for DMSO vs. DEC-treated cells was calculated using the Mann–Whitney test: ****p < 0.0001. Medium (M) to large (L) effect sizes suggest substantial DNA damage, while small (S) effect sizes indicate minimal to no response to DEC treatment. The effect sizes for DEC vs. DMSO treatments were calculated as follows: BxPC-3 (Alkaline: M; Neutral: L), PaTu 8902 (Alkaline: L; Neutral: L), HPAF-II (Alkaline: M; Neutral: S), and CAPAN-1 (Alkaline: M; Neutral: M). No significant increase in TM was observed in indepKRAS cells, with effect sizes calculated as Small (S). Alkaline and neutral comet assays confirmed significant DNA fragmentation indicative of single-strand breaks (SSBs) and double-strand breaks (DSBs) in dKRAS-PDAC cells following DEC treatment. (D) Upper panel: Representative immunoblots showing phosphorylation of CHK1 at Ser345 in the indicated cell lines following DEC (1.25 μM) treatment for 72 hours. Equal amounts of total protein extracts were analyzed with the indicated antibodies. Immunoblot for actin was used as a loading control. Numbers below the actin immunoblots represent the ratio of actin densitometry of each sample relative to DMSO. Lower panel: Histograms represent the quantification of γCHK1(Ser345)/CHK1 densitometry. Histograms represent mean ± SD (represented by error bars) from triplicate experiments. Statistical significance for DMSO vs. DEC-treated cells was calculated using an unpaired t-test: **p < 0.01, ****p < 0.0001, ns: p > 0.05; statistical analysis was performed using GraphPad Prism. (E) Quantification of immunofluorescence analysis of nuclear γCHK1(Ser345) positive foci in dKRAS-PDAC cell models following DEC treatment. Data are presented as median from a representative replicate experiment. At least 50 nuclei were counted for each replicate by image analysis with CellProfiler 4.2.1. Statistical analysis was performed using GraphPad Prism. Statistical significance for DMSO vs. DEC-treated cells was calculated using an unpaired t-test: ***p < 0.001, ****p < 0.0001. (F) Colocalization analysis of γH2AX with γCHK1-positive nuclear foci in dKRAS-PDAC cells following DEC (1.25 μM) treatment for 72 hours. Left panel: Representative images of confocal microscopy analysis of PaTu 8902 and HPAF-II co-stained for γH2AX and with γCHK1. Scale bars, 10 μm. Magnification, x60. Right panel: Dot plots representative of the percentage of colocalized foci. Data are presented as median from a representative replicate experiment. From 20 to 100 nuclei were counted for each replicate by image analysis with CellProfiler 4.2.1. Data analysis was performed using GraphPad Prism. (G) ATR or ATM inhibitors suppress DEC-induced phosphorylation of CHK1 at Ser345 and CHK2 phosphorylation at Thr68 in dKRAS-PDAC cells (PaTu 8902 and HPAF-II). Representative immunoblots of total proteins extracted from indicated cell lines treated with DMSO or DEC (100 nM) for 72 hours. Where indicated, the ATM inhibitor (ATMi) AZD1390 (10 nM) or the ATR inhibitor (ATRi) BAY-1895344 (2 μM) was added 1 hour before harvesting cells. Equal amounts of total protein extracts were analyzed with the indicated antibodies. Immunoblot for actin was used as a loading control. Numbers below the actin immunoblot represent the ratio of actin densitometry of each sample relative to DMSO in each cell line. Quantifications of γCHK1/totalCHK1 and γCHK2/CHK2 are reported in Figure S4.

We further investigated the extent of DNA damage using the comet assay. The comet assay revealed that the tail moment (TM) value, indicative of DNA fragmentation following DNA breaks, was statistically increased in dKRAS-PDAC cells (PaTu 8902, BxPC-3, HPAF-II, and CAPAN-1) following DEC treatment. The effect size for DEC versus DMSO treatment in dKRAS-PDAC cells was calculated to be medium to large, indicating a robust induction of DNA damage (Figure 1C). Conversely, the indepKRAS-PDAC cell lines (PaTu 8988t and KP-4) showed very low or no significant increase in TM values following DEC treatment, with effect sizes calculated to be small (Figure 1C). This differential response underscores the specificity of DEC-induced DNA damage in KRAS-dependent versus KRAS-independent PDAC cells. A significant increase in TM following DEC treatment was observed using both alkaline and neutral comet assay methods, confirming that DEC induces both SSB and DSB DNA damage, as indicated by DDR marker analysis.

Next, we examined the activation of specific DNA damage protein checkpoints. Immunoblot analysis demonstrated that DEC induced the phosphorylation of checkpoint kinase 1 (CHK1) at residue Ser345 in dKRAS-PDAC cell lines, a key residue for kinase activation following DNA damage (Figure 1D). Accordingly, nuclear phospho-Ser345(γ)CHK1-positive foci were detected in dKRAS-PDAC cell models following DEC treatment (Figure 1E), with γCHK1-positive nuclear foci demonstrating colocalization with γH2AX-positive foci (Figure 1F). Inhibiting ATR (Ataxia Telangiectasia and Rad3-related), the key kinase responsible for CHK1 phosphorylation and activation, using a specific ATR inhibitor, completely suppressed DEC-induced phosphorylation of γCHK1 (Figure 1G and Figure S4A). Furthermore, a specific ATM (Ataxia Telangiectasia Mutated) inhibitor completely inhibited DEC-induced phosphorylation at the phospho-Ser68(γ)CHK2 target residue and reduced the extent of DEC-induced CHK1 phosphorylation (Figure 1G and Figure S4B). Overall, these data demonstrate that DEC induces robust DNA damage and DDR in dKRAS-PDAC cells, while indepKRAS cells remain insensitive. The DDR involves ATM/ATR-mediated DNA damage checkpoints through the activation of CHK1 and CHK2 kinases, resulting from the accumulation of both SSB and DSB DNA damage.

### PARP1 activity via PARylation mediates DNA repair and DDR following DEC treatment

Based on the role of PARP1 and PARP activity in both SSB and DSB DNA repair and the previously reported crosstalk with DEC activity, we next investigated the role of PARP1 activation following DEC treatment in dKRAS- and indepKRAS-PDAC cells. Immunoblot analysis of PARP1 extracted from nuclear protein fractions demonstrated that a nanomolar concentration (100 nM) of DEC induced the selective accumulation of PARP1 in the nuclear insoluble (NI), chromatin-bound protein fraction (Figure 2A, upper panels), but not in the nuclear soluble (NS), high-salt extracted protein fraction (Figure 2A, lower panels) in dKRAS-PDAC cells (PaTu 8902 and HPAF-II) but not in indepKRAS cells (PaTu 8988t). DEC increased the number PARP1 nuclear foci and aggregates as revealed by immunofluorescence analysis (Figure 2B) in dKRAS-PDAC cells (PaTu 8902 and HPAF-II) but not in indepKRAS cells (PaTu 8988t).

**Figure 2.**
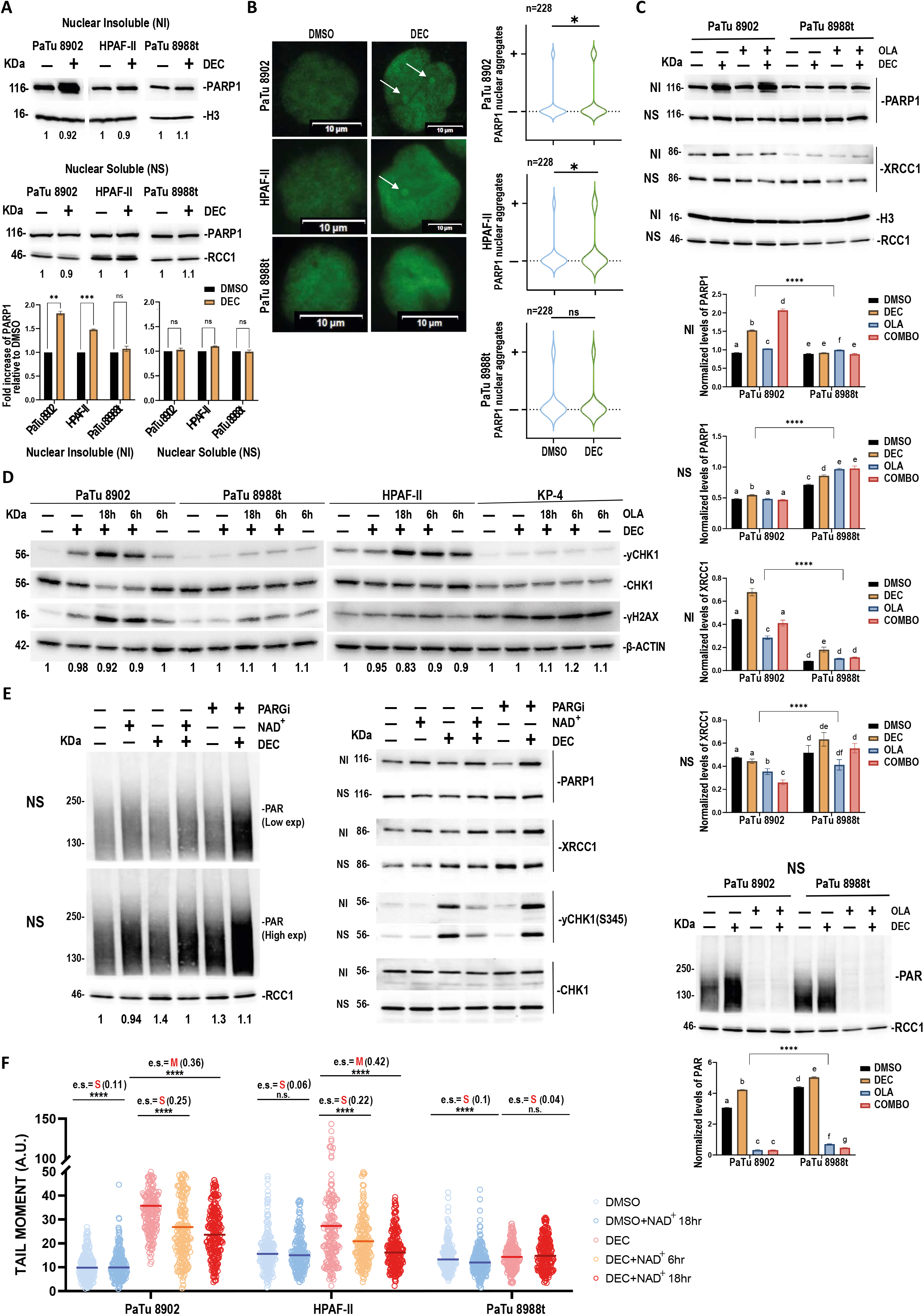
PARP1 activity via PARylation mediates DNA repair and DDR following DEC treatment. (A) Immunoblot analysis of PARP1 in nuclear protein fractions extracted from dKRAS-PDAC cells (PaTu 8902 and HPAF-II) and indepKRAS-PDAC cells (PaTu 8988t) treated with DMSO (vehicle) or decitabine (DEC) at 100 nM for 72 hours. Upper panels show PARP1 accumulation in the chromatin-bound, nuclear insoluble (NI) fraction from dKRAS cells but not from indepKRAS cells. The middle panels display PARP1 levels in the high-salt extracted, nuclear soluble (NS) protein fraction, where no significant changes were observed. Lower panels: Histograms represent the quantification of the PARP1 protein amount over Histone H3 protein used as a loading control for the NI fraction or over RCC1 protein used as a loading control for the NS fraction. Histograms represent mean ± SD from triplicate experiments. Statistical significance for DMSO vs. DEC-treated cells was calculated using an unpaired t-test: **p < 0.01, ***p < 0.001, ns: p > 0.05. (B) Left panels: Representative immunofluorescence images of PARP1 nuclear staining in dKRAS-PDAC cells (PaTu 8902 and HPAF-II) and indepKRAS-PDAC cells (PaTu 8988t) following treatment with DMSO (vehicle) or decitabine (DEC) at 100 nM for 72 hours. Arrows indicate DEC-induced PARP1 protein aggregates (dots) in the nucleus. The nuclei were counterstained with DAPI. Scale bars, 10 μm. Magnification, x60. Right panels: Quantification of PARP1-positive nuclear aggregates as presented in the left panels in dKRAS-PDAC cells (PaTu 8902 and HPAF-II) and indepKRAS-PDAC cells (PaTu 8988t). “+” indicates positive cells presenting at least one PARP1 positive aggregate signal. “-” indicates negative cells, showing no nuclear aggregates of PARP1. Data are presented as mean from a representative replicate experiment. The indicated nuclei (n) were counted for each replicate by image analysis with CellProfiler 4.2.1. Data analysis was performed using GraphPad Prism. Statistical significance for the mean difference of DEC-treated cells vs. DMSO-treated cells was determined using the unpaired two-tailed student t-test. *p < 0.05; ns: p > 0.05. (C) PARylation promotes DEC-induced recruitment of XRCC1 on the chromatin-bound protein fraction. Upper panels: Representative images of immunoblot analysis for PARP1 and XRCC1 in nuclear protein fractions from dKRAS-PDAC cells (PaTu 8902) and indepKRAS-PDAC cells (PaTu 8988t) treated for 72 hours with DMSO, DEC (100 nM), OLA (1.4 μM), or DEC combined with OLA. OLA was added 18 hours before harvesting cells. NI: nuclear insoluble, chromatin-bound protein fraction; NS: nuclear soluble, high-salt extracted protein fraction. Middle panels: Histograms represent the quantification of XRCC1 and PARP1 protein over H3 immunoblot used as loading control for the NI fraction or over RCC1 protein used as loading control for the NS fraction. Histograms represent mean ± SD (represented by error bars) from triplicate experiments. Statistical analyses were performed using GraphPad Prism 8 and R with a 2-way Anova test between pairs of cell lines and One-way ANOVA with multiple comparisons through post-hoc Tukey HSD (Honestly Significant Difference) Test within experimental set of each cell line (same letter in the graph represents no statistical relevance in the difference between groups, different letter represents a significant pvalue (< 0.05) of the comparison between groups). Lower panel: Representative images of immunoblot analysis for PAR (PARylated proteins) in the NS protein fraction, showing the effective inhibition of PARP activity by OLA under the indicated experimental conditions. Histograms represent mean ± SD (represented by error bars) from triplicate experiments Statistical analyses were performed as above described. (D) Activation of DDR markers was significantly enhanced by combined DEC and OLA treatment in dKRAS cells, but not in indepKRAS PDAC cells. Immunoblot analysis of γCHK1(Ser 345) and γH2AX(Ser139) in dKRAS-PDAC cells (PaTu 8902 and HPAF-II) and indepKRAS-PDAC cells (PaTu 8988t and KP-4) treated for 72 hours with DMSO, DEC (100 nM), OLA, or DEC combined with OLA. OLA was added for the indicated time before harvesting cells and used as follows: 1.4 μM for 18 hours; 5 μM for 6 hours. Immunoblot for actin was used as a loading control. Numbers below the actin immunoblots represent the ratio of actin densitometry of each sample relative to DMSO. The quantifications of γCHK1 and γH2AX are reported in Figure S5A. (E) Left panels: Enhancing PARP1 activity by Nicotinamide adenine dinucleotide (NAD+) supplementation. dKRAS-PDAC cells (PaTu 8902) were treated with DMSO or DEC (100 nM) for 72 hours and supplemented with NAD+ (0.5 mM) for 3 hours before harvesting the cells. Immunoblot analysis (low and high exposures) of the nuclear soluble protein fraction showed increased protein PARylation, indicating enhanced PARP activity following NAD+ supplementation. As a positive control, the poly(ADP ribose) glycohydrolase (PARG) inhibitor PDD00017273 (1 μM) was added 1 hour before harvesting cells. Adding PARGi resulted in the accumulation of PARylated proteins. Quantifications of immunoblots are reported in Figure S5B. Right panels: Enhancing PARP1 activity by NAD+ supplementation in DEC-treated cells increased the amount of chromatin-bound, nuclear insoluble PARP1 and XRCC1, and reduced the extent of DEC-induced γCHK1. Left panels: Representative images of immunoblot analysis for PARP1, XRCC1, γCHK1, and CHK1 in nuclear fractions from experiments performed as in (E). NI: nuclear insoluble, chromatin-bound fraction; NS: nuclear soluble, high-salt extracted protein fraction. Quantifications of immunoblots are reported in Figure S5C-E. (F) Enhancing PARP1 activity by NAD+ supplementation significantly reduced DEC-induced DNA fragmentation in dKRAS cells, indicating enhanced DNA repair by PARylation. Quantification of comet tail moment (TM) values from neutral comet assays of dKRAS-PDAC cells (PaTu 8902, HPAF-II) and indepKRAS-PDAC cells (PaTu 8988t) treated with DMSO, DEC (100 nM), and supplemented, where indicated, with NAD+ (0.5 mM) for 6 or 18 hours before harvesting the cells for comet assays. Data are presented as median from a representative replicate experiment. At least 200 nuclei were counted for each replicate. To quantify the magnitude of treatment-induced DNA damage among groups, the effect size (e.s.) was estimated by the Wilcoxon test by three to seven biological replicates. The Rstatix package was implemented for the calculation of the effect size. Statistical significance among the indicated comparison groups was calculated using the Dunn’s test. Statistical significance: ****, p < 0.0001. Medium (M) to large (L) effect sizes suggest substantial DNA damage, while Small (S) effect sizes indicate minimal to no response to treatment. Effect sizes (e.s): S = Small; M = Medium; L = Large. The calculated effect sizes indicate a robust induction of DNA damage by DEC that could be in part rescued by NAD+ supplementation. No significant increase in TM was observed in indepKRAS cells (PaTu 8988t) following DEC or NAD+ treatments, with effect sizes calculated as Small (S).

To address the mechanistic role of PARP1 and PARP activity in response to DEC in PDAC, we first monitored molecular effects following acute PARP activity inhibition using the FDA-approved PARP inhibitor Olaparib (OLA) in cells pre-treated for 72 hours with a nanomolar concentration of DEC (100 nM). Adding OLA alone for 18 hours before harvesting cells further increased PARP1 accumulation in the nuclear insoluble (NI), chromatin-bound fraction in DEC-treated PaTu 8902 cells. Drug treatments did not increase levels of PARP1 in the nuclear soluble (NS), high-salt extracted protein fraction (Figure 2C). In contrast, in indepKRAS cells (PaTu 8988t), we did not observe accumulation of PARP1 in the NI protein fraction. Since PARylation of PARP1 is known to recruit repair effectors such as XRCC1 to the damaged site for repair, we monitored the amount of XRCC1 in the nuclear fractions. Indeed, DEC treatment increased the amount of XRCC1 in NI fraction in PaTu 8902 cells. Treatment with OLA was sufficient to fully inhibit DEC-induced XRCC1 accumulation within the NI, chromatin-bound protein fraction. IndepKRAS cells (PaTu 8988t) showed a low accumulation of XRCC1 following DEC in the NI fraction, which was also inhibited by OLA treatment (Figure 2C).

To investigate the functional consequence of the inhibition of PARP activity in DEC-treated cells, we monitored the activation of DDR markers. Immunoblot analyses showed that acute treatment with OLA following DEC treatment was sufficient to enhance the activation of DDR markers such as γCHK1 and γH2AX in dKRAS-PDAC cells (PaTu 8902 and HPAF-II), compared to DEC or OLA treatment alone. In contrast, in indepKRAS-PDAC cells (PaTu 8988t and KP-4), DEC was ineffective, and adding OLA did not enhance the extent of DDR marker activation compared to the effect of treating with OLA alone (Figure 2D and Figure S5A).

We next tested the effects of promoting PARP activity. To this aim, we supplemented DEC-treated cells with Nicotinamide adenine dinucleotide (NAD^+^), a cofactor necessary for PARP enzymatic activity. The supplementation of NAD+ was effective in increasing PARP1 activity, as measured by the specific analysis of total protein PARylation by immunoblot (Figure 2E, Left panels, and Figure S5B). This effect was mirrored by treating cells with a Poly (ADP-ribose) glycohydrolase inhibitor (PARGi), which, by inhibiting the de-PARylation of PARylated proteins, served as a positive control for detecting protein PARylation enhancement. Immunoblot studies demonstrated that enhancing PARylation either by NAD+ or PARGi treatments was sufficient to increase the amount of PARP1 in the chromatin-bound, nuclear insoluble (NI) protein fraction, but not in the soluble nuclear fraction, following DEC treatment (Figure 2E, Right panels, and Figure S5C). These results were consistent with the well-established role of PARP1 PARylation in stabilizing PARP1 recruitment on damaged chromatin to recruit DNA repair complexes. Accordingly, in DEC-treated cells, NAD^+^ supplementation also increased the amount of XRCC1 in the chromatin-bound fraction resulting in a parallel reduction of XRCC1 protein amount in the soluble nuclear fraction of DEC-treated cells (Figure 2E, Right panels, and Figure S5D). This effect was also mirrored by the PARGi. Furthermore, the enhancement of PARP1 activity by NAD+ was sufficient to reduce the extent of DEC-induced γCHK1 activation (Figure 2E, Right panels, and Figure S5E), indicating a reduction in the DNA damage response. Interestingly, PARGi did not reduce γCHK1 levels in DEC-treated samples, suggesting persistent DNA damage. This result is expected and is well-consistent with the established role of PARylation/de-PARylation cycles of PARP1 required for proper repair mechanisms following DNA damage, where PARG activity was demonstrated to be essential to support the dynamic assembly/disassembly of multienzyme complexes on damaged DNA for repair.

The enhancement of PARP1 activity by NAD^+^ reduced the extent of γCHK1 activation (Figure 2E, Right panels), suggesting reduced DNA damage. To confirm this, we analyzed the extent of DEC-induced DNA damage and fragmentation using the comet assay. Indeed, the effect size analysis of the tail moment (TM) indicated a robust induction of DNA damage by DEC in dKRAS-PDAC cells (PaTu 8902 and HPAF-II). This damage was partially reduced by NAD+ supplementation, with a calculated effect size of moderate (M) when comparing DEC-treated cells to DEC-treated cells supplemented with NAD+ (Figure 2F).

Overall, these results demonstrated that PARP1 activity played a mechanistic role in the DDR and DNA damage repair induced by DEC in KRAS-dependent PDAC. The acute inhibition of PARP activity by OLA enhances DEC-induced DNA damage, highlighting the potential for combined therapeutic strategies targeting PARP activity in KRAS-dependent PDAC.

### Synergistic combination of low-dose DEC with OLA in dKRAS-PDAC cells

The above results prompted us to investigate whether OLA might enhance the antitumor activity of DEC even when used at a low dose against dKRAS-PDAC. Due to the well-established efficacy of PARPi such as OLA in the context of *BRCA1/2* mutated tumors, we considered the effects of drug treatment in both dKRAS-/*BRCA1/2*^*wild-type*^ PDAC cell lines (PaTu 8902, HPAF-II) and dKRAS-/*BRCA2*^*mut*^ PDAC cells (CAPAN-1). We utilized the Combenefit software to analyze synergy, additivity, or antagonism between the OLA and DEC drug combination in a seven-day experiment. A matrix model (HAS) of combination analysis showed that OLA and DEC were highly synergistic or additive in dKRAS-/*BRCA1/2*^*wild-type*^ PDAC cell lines across the entire range of tested concentrations (Figure 3A). Notably, DEC and OLA were highly synergistic in dKRAS-/*BRCA2*^*mut*^ PDAC cells (CAPAN-1) across all tested concentrations. The synergistic effect was further demonstrated through the Calcusyn method in a seven-day experiment. Figure S7 shows plots of the combination index calculated for the interaction between the two drugs. Interestingly, both models predicted that in dKRAS-/*BRCA1/2*^*wild-type*^ PDAC, the effect moved from synergistic to additive when both drugs were used at higher concentration ranges, supporting the advantage of low-dosage drug combinations, which would be beneficial in a clinical context. Additionally, both models predicted that in dKRAS-/*BRCA2*^*mut*^ PDAC cells (CAPAN-1), the combination was synergistic across all tested concentrations.

**Figure 3.**
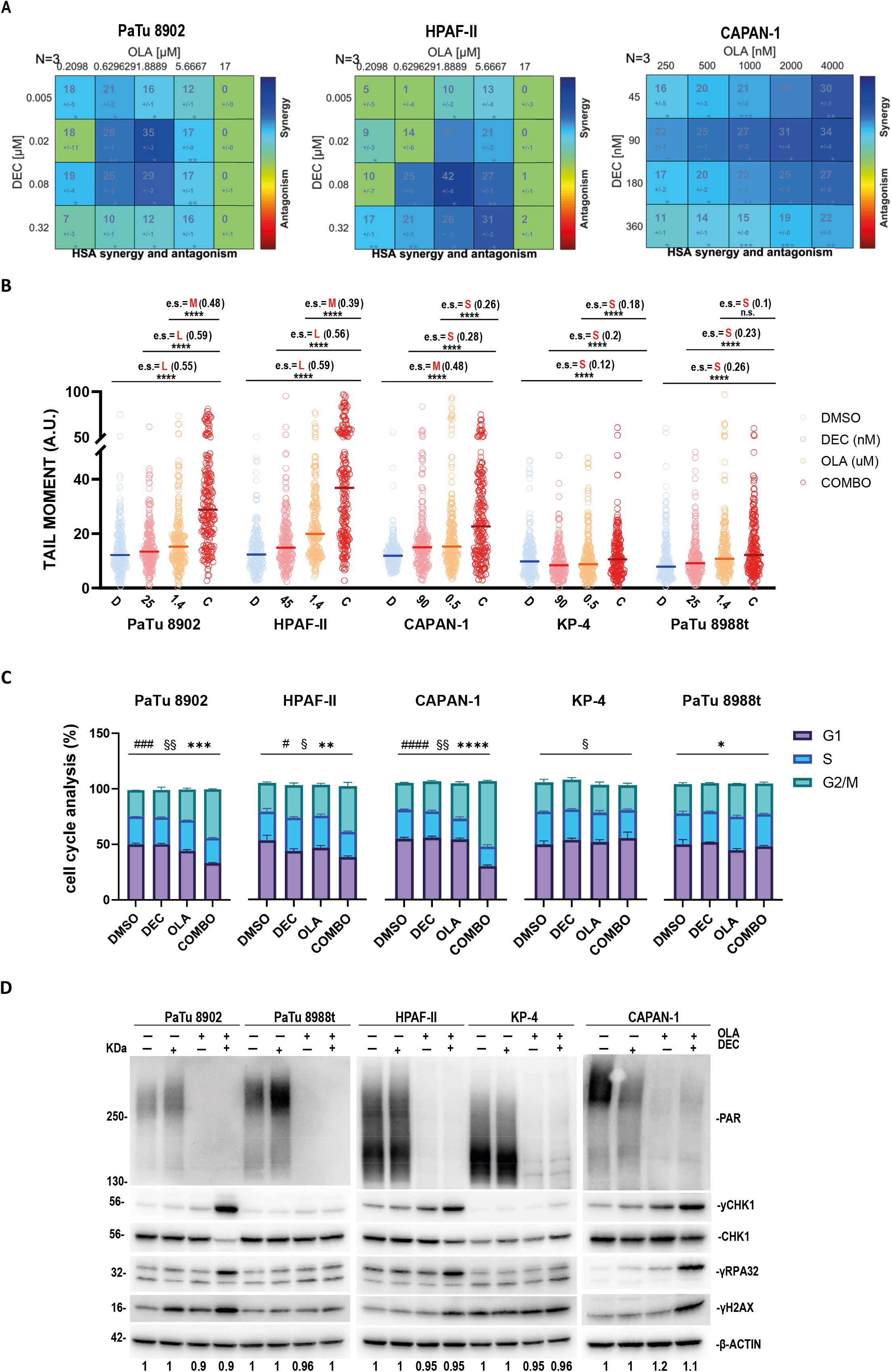
Synergistic combination of low-dose DEC with OLA in dKRAS-PDAC cells. (A) DEC synergizes with OLA. The drug matrix Heatmap, presented in a 4×5 grid, illustrates the HAS model index as analyzed by Combenefit methods for pharmacological combination index in dKRAS-/*BRCA1/2*^*wild-type*^ PDAC cell lines (PaTu 8902, HPAF-II) and dKRAS-/*BRCA2*^*mut*^ PDAC cells (CAPAN-1). Drug combination doses that exhibit synergy appear blue on the heatmap. The indicated cell lines were plated and grown for 6 days with the specified doses of DEC and OLA. Cell viability was then assessed using the ATP-based CellTiter-Glo assay. Data analysis was conducted using Combenefit software. n=3. Drug concentration ranges were determined based on in vitro calculated half-maximal inhibitory concentration (IC_50_) values in each cell line over a seven-day experiment period (Figure S6). (B) Quantification of comet tail moment (TM) was performed in dKRAS-PDAC cells (PaTu 8902, HPAF-II, and CAPAN-1) and indepKRAS-PDAC cells (PaTu 8988t and KP-4) following 72-hour treatment with DMSO, low-dose DEC, OLA, or simultaneous DEC plus OLA (COMBO) at the indicated drug concentrations selected based on Combination Index results (Figure 3A). Data are presented as the median from a representative replicate experiment of the neutral comet assay. At least 200 nuclei were counted for each replicate. To quantify the magnitude of treatment-induced DNA damage among groups, the effect size (e.s.) was estimated using the Wilcoxon test with three biological replicates. The Rstatix package was utilized for the calculation of the effect size. Statistical significance among the indicated comparison groups was determined using the Dunn’s test. Statistical significance is denoted by ****, indicating p < 0.0001. Medium (M) to large (L) effect sizes suggest substantial DNA damage, while small (S) effect sizes indicate minimal to no response to treatment. The calculated effect sizes indicate a robust induction of DNA damage by COMBO compared to single drugs in dKRAS-cells. No significant increase in TM was observed in indepKRAS cells following COMBO treatment, with effect sizes calculated as small (S). Results of the alkaline comet assay are reported in Figure S8. C) Low-dose Decitabine in combination with OLA induced a robust cell-cycle block at the G2/M phases of dKRAS-PDAC cells but not of indep-KRAS cells. Cells were treated for 72 hours with DMSO, low-dose DEC, OLA, or simultaneous DEC plus OLA (COMBO) at drug concentrations as indicated in (B) and then assayed by FACS analysis of propidium iodide-stained cells. Histograms show mean ± SD (n=3). Statistical significance among treatments was calculated using a 1-way ANOVA. # refers to G1-phase; § refers to S-phase; * refers to G2/M-phase of the cell cycle. One symbol: p ≤ 0.05; two symbols: p ≤ 0.01; three symbols: p ≤ 0.001; four symbols: p ≤ 0.0001. (D) DDR markers were activated by COMBO treatment, compared to low-dose DEC or OLA alone, in dKRAS-/*BRCA1/2*^*wild-type*^ PDAC cell lines (PaTu 8902, HPAF-II) and dKRAS-/*BRCA2*^*mut*^ PDAC cells (CAPAN-1) but not in indepKRAS-/*BRCA1/2*^*wild-type*^ PDAC cell lines (PaTu 8988t and KP-4). Immunoblot analysis of γCHK1(Ser 345), γRPA32(Ser4/Ser8), and γH2AX(Ser139) in the indicated cell lines treated for 72 hours with DMSO, DEC, OLA, or COMBO at drug concentrations as in (B). Immunoblot for actin was used as a loading control. Numbers below the actin immunoblots represent the ratio of actin densitometry of each sample relative to DMSO. The quantifications of γCHK1(Ser345), γRPA32(Ser4/Ser8), and γH2AX(Ser139) activation are reported in Supplementary Figure S9.

Based on combination index results, we tested the cytotoxic effects of an even lower nanomolar concentration of DEC in combination with OLA (hereafter referred to as “COMBO”) by simultaneously co-treating cells with both drugs in functional assays in vitro. We first monitored the genotoxic effect of COMBO by comet assay (Figure 3B). Results demonstrated that the tail moment significantly increased in dKRAS-/*BRCA1/2*^*wild-type*^ PDAC cell lines (PaTu 8902, HPAF-II) and dKRAS-/*BRCA2*^*mut*^ PDAC cells (CAPAN-1) treated for 72 hours with COMBO. Indeed, the calculated effect size of the tail moment in COMBO-treated cell lines was large (L) when compared to untreated cells or single-drug treatments alone. The increase in the tail moment following COMBO treatment was observed by both alkaline and neutral comet assay methods. Conversely, the indepKRAS-/ *BRCA1/2*^*wild-type*^ PDAC cell lines (PaTu 8988t and KP-4) showed very low or no significant increase in tail moment values following COMBO treatment, with effect sizes calculated to be small compared to the vehicle as well as to single agents alone (Figure 3B and S8).

COMBO, but not the single treatments, induced a significant G2/M-phase arrest of the cell cycle in dKRAS-/*BRCA1/2*^*wild-type*^ PDAC cell lines (PaTu 8902 and HPAF-II) and dKRAS-/*BRCA2*^*mut*^ PDAC cells (CAPAN-1) (Figure 3C). This strong and significant G2/M-phase arrest was not observed in indepKRAS-/ *BRCA1/2*^*wild-type*^ PDAC cell lines (PaTu 8988t and KP-4). Immunoblot analyses showed that COMBO treatment, while effectively reducing PARylation (Figure 3D and Figure S9A), enhanced the activation of DDR markers such as γCHK1, γRPA, and γH2AX in dKRAS-/*BRCA1/2* wild-type PDAC cell lines and dKRAS-/*BRCA2* mutant PDAC cells compared to DEC or OLA alone, or to the vehicle DMSO (Figure 3D and Figure S8B-D).In contrast, COMBO was ineffective in increasing DDR marker activation in indepKRAS-/ *BRCA1/2*^*wild-type*^ PDAC cell lines compared to OLA or the vehicle alone (Figure 3D and Figure S8B-D). Overall, in vitro studies demonstrated the antiproliferative efficacy of combining very low-dose DEC with OLA in both dKRAS-/*BRCA1/2*^*wild-type*^ and dKRAS-/*BRCA2*^*mut*^ PDAC cells.

### Synergistic antitumor activity of low-dose decitabine with OLA in dKRAS-/*BRCA1/2*^*Wild-Type*^ tumor xenograft models

The antitumor activity of the combination treatment was evaluated next. We implanted dKRAS-/*BRCA1/2*^*wild-type*^ PDAC tumor cells (HPAF-II) or indepKRAS-/*BRCA1/2*^*wild-type*^ PDAC tumor cells (KP-4 and PaTu 8988t) subcutaneously in NSG-immune-deficient mice. Xenograft tumor models were then subjected to treatment with either vehicle, low-dose DEC (0.2 mg/kg), OLA, or the combination treatment (COMBO). In comparison to higher doses of DEC, low-dose DEC treatment was well-tolerated and exhibited a better safety profile in experimental models (Figure S10A). Results indicated that COMBO treatment led to a significantly synergistic antitumor effect, reducing the growth of dKRAS-/*BRCA1/2*^*wild-type*^ PDAC xenograft tumors compared to single treatments alone (Figures 4A-B, S10B). There were no discernible effects in indepKRAS-/*BRCA1/2*^*wild-type*^ PDAC xenograft tumors following combined treatment (Figures 4C-D and 4E-F, S10C-D) when compared to OLA treatment alone. Furthermore, while displaying enhanced antitumor activity, COMBO treatment was well tolerated in vivo, as no significant changes in body weight percentage were observed in COMBO-treated mice compared to those receiving single treatment or vehicle alone (Figure S10F-I).

**Figure 4.**
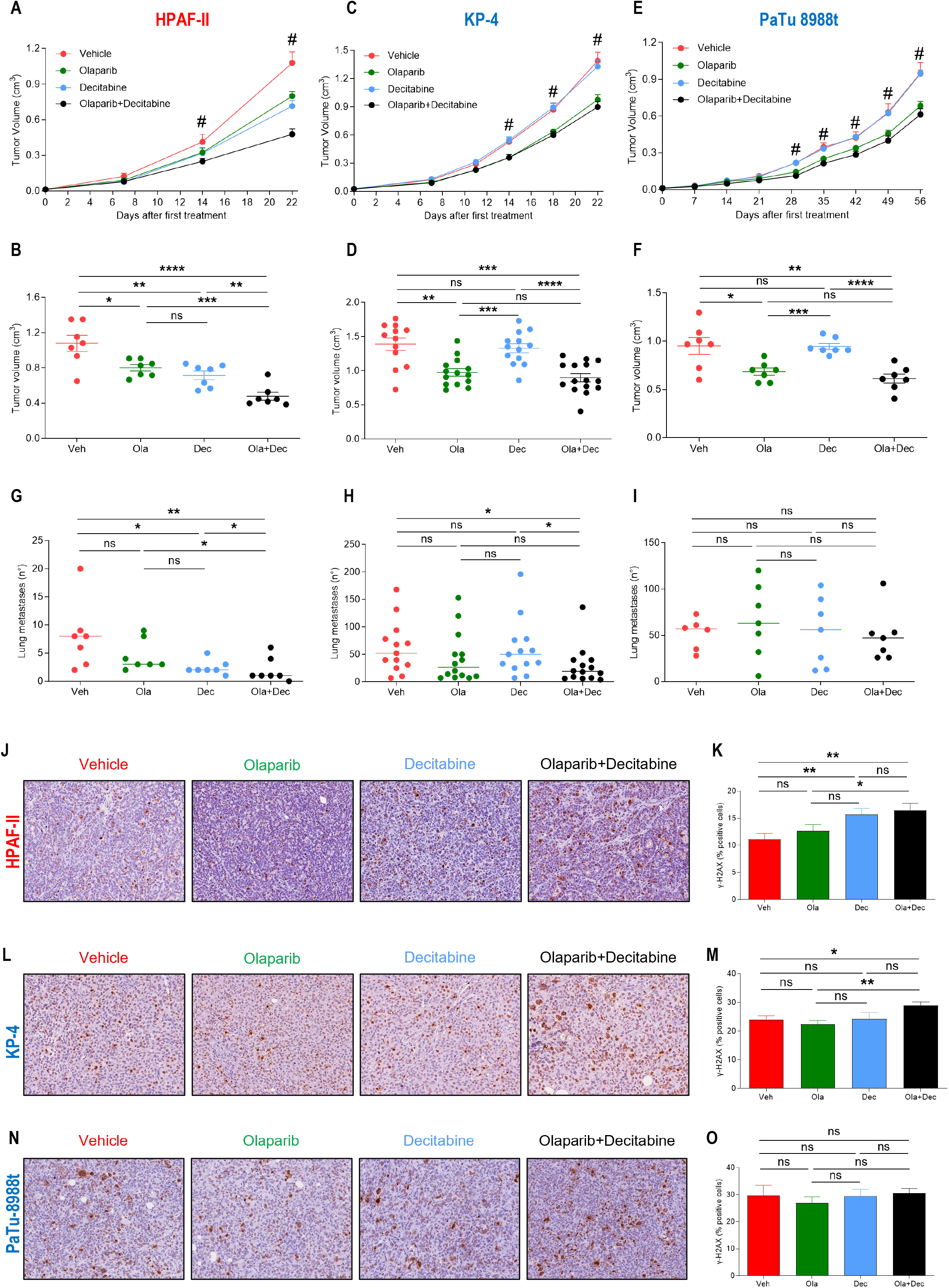
Synergistic antitumor activity of low-dose decitabine with OLA in dKRAS-/*BRCA1/2*^Wild-Type^ tumor xenograft models. (A) Tumor growth kinetics and tumor volume at day 22 (B) of mice injected subcutaneously with the HPAF-II cell line and treated with Vehicle (red line and point), Olaparib (green line and point), Decitabine (blue line and point), and a combination of Olaparib and Decitabine (black line and point). Tumor growth (C) and tumor volume at day 22 (D) of mice injected subcutaneously with the KP-4 cell line, following different treatments; tumor growth kinetics (E) and tumor volume at day 56 (F) of mice injected subcutaneously with the PaTu 8988t cell line and treated with the different drugs. The number of lung metastases of mice injected with HPAF-II (G), KP-4 (H), and PaTu 8988t (I) cell lines. Representative images of immunohistochemical staining of γH2AX marker in sections prepared from HPAF-II tumors (J), KP-4 tumors (L), and PaTu-8988t tumors (N) from each treatment group at the end of the experiments. Histograms show the quantitative evaluation of γH2AX expressed as a percentage of positive cells in the tumor area of HPAF-II tumors (K), KP-4 tumors (M), and PaTu 8988t tumors (O). Data are represented as mean ± SEM (A, B, C, D, E, F, K, M, O) or median (G, H, I). Differences in tumor growth kinetics were evaluated using a two-way ANOVA test analysis, while differences in tumor volumes and the percentage of γH2AX-positive cells were evaluated using unpaired t-test analysis; differences in the number of lung metastases were evaluated using Mann–Whitney test analysis. *, p < 0.05; **, p < 0.01; ***, p < 0.001; ****, p < 0.0001; ns, not significant.

Analysis of hematoxylin-eosin-stained lung tissue sections revealed that COMBO treatment exhibited greater antimetastatic effects compared to single drug treatments, as evidenced by a reduction in the number of spontaneous lung metastases generated by dKRAS-/*BRCA1/2*^*wild-type*^ PDAC xenograft tumors (Figure 4G). Conversely, COMBO treatment showed no significant effects on reducing the number of spontaneous metastases originating from indepKRAS-/*BRCA1/2*^*wild-type*^ PDAC, with effects limited to OLA alone, but unresponsive to DEC and COMBO (Figure 4H-I).

To investigate whether COMBO induced DNA damage in vivo, immunohistochemistry (IHC) studies were conducted. Consistent with in vitro experiments, COMBO treatment led to increased staining of γH2AX-positive nuclei in dKRAS-/*BRCA1/2*^*wild-type*^ PDAC xenograft tumors (Figure 4J-K). However, IHC analysis of the same markers in indepKRAS-/*BRCA1/2*^*wild-type*^ PDAC treated with COMBO showed no significant differences compared to DMSO (vehicle)-treated tumors, with effects limited to those elicited by OLA alone (Figure 4L-O).

### Synergistic antitumor activity of low-dose decitabine with OLA in dKRAS-/*BRCA2*^*mut*^ PDAC

The results of the in vitro combination index assay revealed significant synergy in dKRAS/g*BRCA2*mut PDAC cells (CAPAN-1), as COMBO treatment exhibited synergy across all tested drug concentrations. This suggests exceptional efficacy of COMBO in the subgroup of dKRAS-/*BRCA2*^*mut*^ PDAC, which would already be eligible for OLA treatment. Xenograft tumors derived from CAPAN-1 cells further confirmed that the COMBO treatment displayed higher antitumor activity compared to single treatments (Figures 5A-B, S10E). Notably, the combination treatment exhibited significantly greater efficacy against metastasis growth, achieving complete inhibition compared to DEC or OLA alone (Figure 5C). As expected, COMBO treatment led to increased staining of γH2AX-positive nuclei in xenograft tumors (Figure 5D-E). These findings underscore the high synergistic activity of the combination therapy in the context of dKRAS-PDAC tumors already candidates for OLA therapy, where COMBO showed a strong antimetastatic effect, leading to complete metastasis growth inhibition.

**Figure 5.**
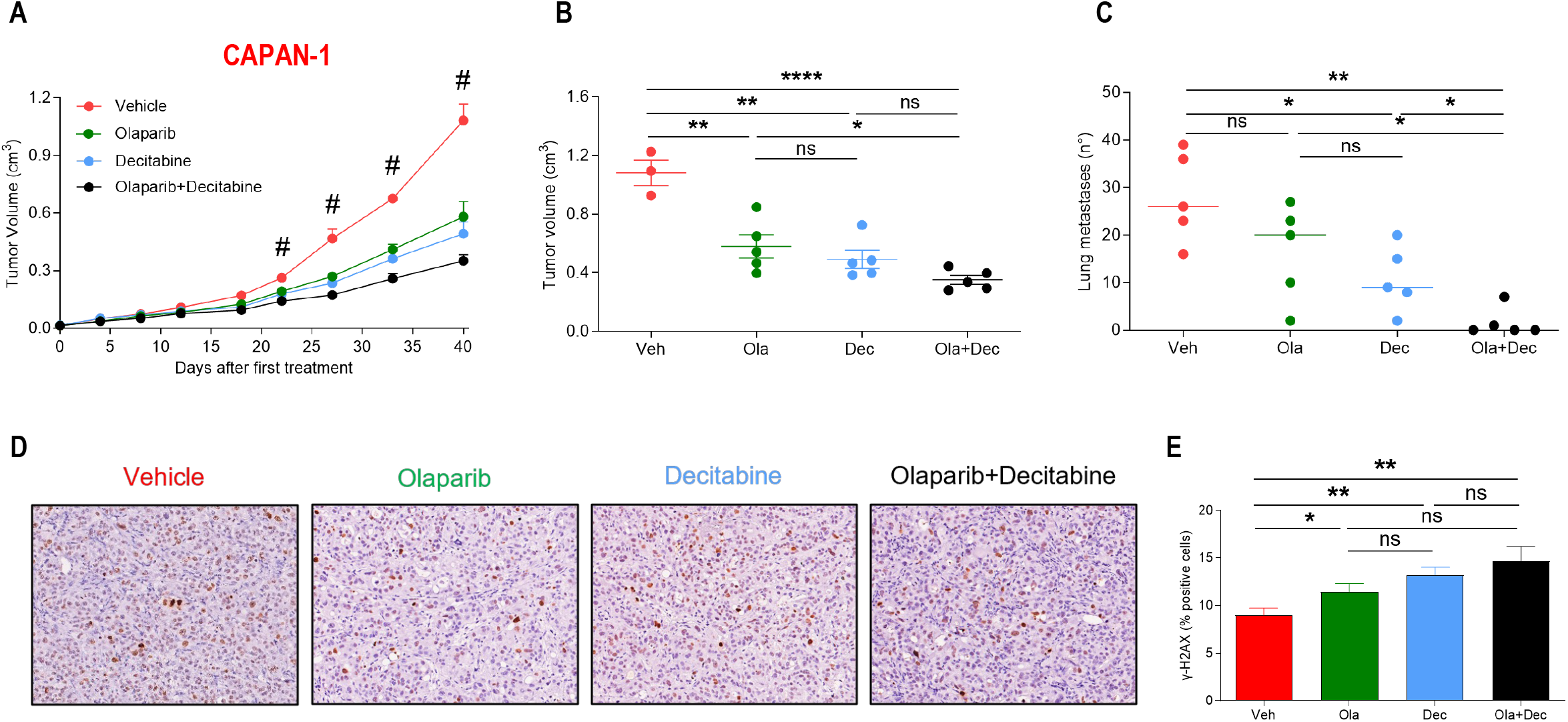
Synergistic antitumor activity of low-dose decitabine with OLA in dKRAS-/*BRCA2*^mut^ PDAC. (A) Tumor growth kinetics and tumor volume at day 40 (B), and number of lung metastases (C) in mice injected subcutaneously with the CAPAN-1 cell line and treated with Vehicle (red line and point), Olaparib (green line and point), Decitabine (blue line and point), and a combination of Olaparib and Decitabine (black line and point). Representative images of immunohistochemical staining of the γH2AX marker in tumor sections (D), and quantitative evaluation of γH2AX expression as a percentage of positive cells in the tumor area (E). Data are represented as mean ± SEM (A, B, E) or median (C). Differences in tumor growth kinetics were evaluated using a two-way ANOVA test analysis, while differences in tumor volume and the percentage of γH2AX-positive cells were evaluated using unpaired t-test analysis; differences in the number of lung metastases were evaluated using Mann–Whitney test analysis. *, p < 0.05; **, p < 0.01; ***, p < 0.001; ****, p < 0.0001; ns, not significant.

### KRAS dependency and response to COMBO therapy are not directly associated with Homologous Recombination Deficiency (HRD) biomarkers

To further investigate the predictive value of KRAS dependency for combination treatment with OLA and DEC, we assessed whether the selective efficacy of COMBO treatment could be predicted based on established molecular biomarkers of OLA’s antitumor efficacy, such as HRD or BRCAness-like conditions. This analysis is crucial to determine if KRAS dependency serves as a unique, independent molecular biomarker for predicting response to COMBO in PDAC. To this aim, we analyzed the genomic mutational profiles of homologous recombination (HR)-related genes in dKRAS- and indepKRAS-PDAC cell lines under investigation in this study using publicly available mutational data from the COSMIC and cBioPortal databases. We searched for both gene mutations and copy number alterations (CNAs) in genes commonly involved in DNA damage response (DDR) and homologous recombination, whose mutations are known to be associated with increased OLA cytotoxicity (Figure 6A). Our database inspection revealed that, apart from CAPAN-1 cells harboring germline *BRCA2*, *ATR*, and *FANC1* mutations and PaTu 8988t cells carrying a nonsense mutation in *MRE11A*, no additional functionally relevant mutations were found in the selected HR-related genes. Furthermore, no CNAs were detected in HR-related genes in the investigated cell lines, except for *BRCA1*, *FANC1*, and *MLH1* genes in the CAPAN-1 cell line (Figure 6A). Additionally, we analyzed microsatellite stability (MS) using reported and validated annotations for the overall CCLE panel, and results demonstrated that all investigated cell models exhibited stable microsatellite (MSS) scores (Figure 6A).

**Figure 6.**
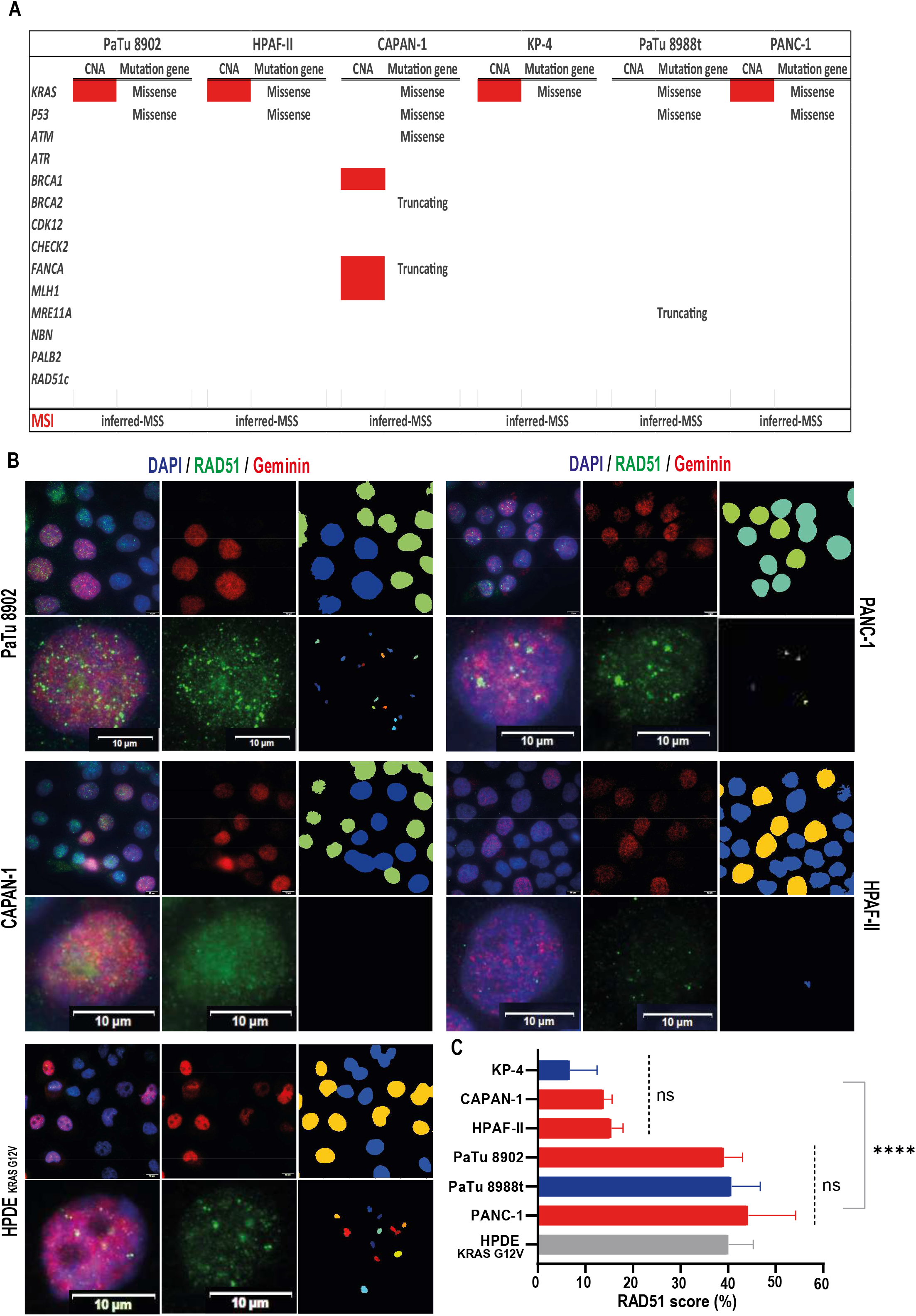
KRAS dependency and response to COMBO therapy are not directly associated with Homologous Recombination Deficiency (HRD) biomarkers. (A) Annotation of genomic mutations and gene copy number alterations (CNAs), and inferred Microsatellite (MS) stability status of a panel of HR-related genes in the indicated PDAC cell lines obtained through inspection of the COSMIC and the cBioPortal databases. A red box indicates > 2 gene copies. Where indicated, the type of retrieved mutations is annotated. Inferred-MSS: Microsatellite stability, according to previous reports. (B) Representative immunofluorescence analysis of nuclear RAD51 foci-/Geminin-positive nuclei in a panel of PDAC cell lines for the determination of the “RAD51 score” as a functional biomarker of HR-proficient cells. The nuclei were counterstained with DAPI. Scale bars, 10μm. Magnification, x60. i) Representative image of merged nuclear signals for RAD51/DAPI/Geminin; ii) representative magnification of a single cell as stained in i); iii) representative images of Geminin-positive nuclei; iv) representative magnification of RAD51 nuclear foci; v) representative image obtained through computational segmentation by CellProfiler 4.2.1 software for Geminin-positive cells and automated cell counting; vi) representative image obtained through computational segmentation by CellProfiler 4.2.1 software for nuclear RAD51-positive foci for an automated cell counting. PANC-1 is a well-established HR-proficient PDAC cell line; HPDE-KRAS^*G12V*^: Human Pancreatic Ductal Epithelial cells, stably transfected with oncogenic KRAS^*G12V*^, here referred to as a control of an HR-proficient cellular model. (C) Histogram showing RAD51 scores for PDAC cell lines. Blue bars: indepKRAS-cell lines; red bars: dKRAS-cell lines. The “RAD51 score” was calculated as the percentage of RAD51-positive nuclei (≥ 3 nuclear RAD51 foci) over Geminin-positive nuclei. At least 200 nuclei were counted for each replicate by image analysis with CellProfiler 4.2.1 as represented in B. Data are presented as mean ± SD. Statistical analysis was performed using GraphPad Prism. ns: no statistically significant difference within the RAD51 score high group and the RAD51 score low group was calculated by the Kruskal-Wallis test (dotted lines). Statistical significance among the RAD51 score high group versus the RAD51 score low group of cell lines was calculated using a two-way ANOVA test, ****, p < 0.0001 (solid line).

We further investigated the status of the so-called “RAD51 score” as a potential cellular biomarker predicting HR functional status, since the RAD51 score has gained interest as a predictor of OLA sensitivity, as detailed in the Introduction. We estimated the RAD51 score in the panel of PDAC cell lines used for this investigation by immunofluorescence analysis and calculated it as the percentage of RAD51 nuclear foci-positive/Geminin-positive cells. The RAD51 score was demonstrated to be low in the *BRCA2*-mutated cells, while being high in the HR-proficient cells PANC-1 or in non-transformed human pancreatic epithelial cells (HPDE) carrying the oncogenic *KRAS*^*G12V*^ mutation (HPDE-KRAS^*G12V*^), used as a control. The RAD51 score analysis identified a subgroup of cell lines with a high RAD51 score (average score: 41.1% ± 2.2) and a subgroup of cell lines with a low RAD51 score (average score: 12% ± 4.6). Results showed no association with KRAS dependency status, as KRAS-dependent models belonged to both the high and low RAD51 groups.

Overall, our findings suggest that KRAS dependency biomarkers, as determined by the L and S scores in PDAC cell lines, are not strictly associated with known HRD biomarkers such as gene mutations in HR-related genes and microsatellite instability. Therefore, KRAS dependency scores may represent a novel, independent biomarker sufficient to predict response to COMBO treatment in PDAC. However, based on results from the CAPAN-1 and HPAF-II models (Figures 3-5), tumor cells carrying both biomarkers, i.e., KRAS dependency with a *BRCA2* mutation or low RAD51 score, demonstrated high responsiveness, suggesting that combining dKRAS-dependency scores with HRD biomarkers may be further explored as predictors of drug response, potentially useful for enhancing personalized therapies involving DEC and PARPi.

## DISCUSSION

Pancreatic ductal adenocarcinoma (PDAC) remains one of the most lethal solid tumors, with limited treatment options and poor prognosis. The emergence of personalized medicine has led to a growing interest in identifying targeted therapies for PDAC subtypes, particularly those driven by specific genetic alterations such as KRAS mutations. In this study, we investigated the therapeutic potential of combining low-dose decitabine (DEC) with olaparib (OLA) in PDAC, focusing on the subset of tumors exhibiting KRAS dependency. Our findings provide compelling evidence for the synergistic antitumor activity of this combination therapy in KRAS-dependent PDAC models. Notably, the synergistic effect was independent of BRCA mutations, suggesting broader applicability beyond *BRCA*-mutated tumors.

The rationale for combining DEC and OLA stems from their complementary mechanisms of action. DEC, a DNA methyltransferase inhibitor, induces DNA damage and sensitizes tumor cells to DNA damage repair inhibitors such as OLA. Preclinical studies here presented demonstrated that this combination results in enhanced cytotoxicity and antimetastatic effects in KRAS-dependent PDAC models, offering a promising therapeutic strategy for this challenging disease utilizing low-dose, well-tolerated drug combination regimens of DEC plus OLA.

Despite over fifty clinical trials testing the efficacy of decitabine across various cancers, including breast, ovarian, colon, lung, pancreatic cancers, and sarcoma, its toxicity remains a key limiting factor for assessing its antitumor efficacy. FDA-approved regimens are associated with a high rate of hematological toxicity in approximately 30% of cases, including grade 3/4 febrile neutropenia. Importantly, DEC’s adverse events may be mitigated by different treatment schemes (such as metronomic dosing) and by reducing dosages ^24,25^ raising questions about the impact of these changes on its antitumor efficacy. In this context, a combination of synergistic drugs might provide the key. Drug combinations in oncology can improve patient prognosis when a synergistic therapeutic efficacy is achieved. Additionally, synergistic drug combinations at low doses increase the likelihood of achieving drug efficacy at doses that can be readily attained in clinical settings compared to those tested in preclinical studies. Even if the drug synergism observed in preclinical settings results in drug additivity in clinical settings, additivity still represents significant gains in improving therapeutic outcomes ^26^. Furthermore, for the COMBO treatment analyzed here, the different hematological toxicity profiles of each drug (i.e., neutropenia for decitabine and anemia for olaparib) are expected to provide more manageable severe adverse events.

From a mechanistic point of view, the combination of OLA and DEC was linked to DEC’s ability to induce single-strand breaks (SSB) and double-strand breaks (DSB) DNA damage, highlighting its broader potential as an anticancer agent beyond its established role in gene expression regulation. Several studies have demonstrated that DEC can induce DNA damage in HeLa, colon cancer, and multiple myeloma cells ^15,16^. The base excision repair (BER) pathway has been implicated, at least in part, in the removal of aberrant bases generated by DEC ^17^. The antitumor activity of the decitabine and PARP inhibitors (PARPi) combination has substantial mechanistic evidence in tumors other than PDAC. Increased cytotoxic effects of PARP inhibitors with DNA demethylating agents have been reported in unfavorable AML subtypes and BRCA wild-type breast cancer cells ^23^, as well as in breast and ovary cancer [Pullam, 2018] and NSCLC, where DNA methyltransferase inhibitors induced a BRCAness phenotype able to sensitize NSCLC to PARP inhibitor and ionizing radiation ^27,28^ The cytotoxic activity of covalent PARP1/DNA adducts was also observed in non-transformed human cells following decitabine treatment ^29^. Recent studies have further implicated PARP1 activity in DSB repair, where PARP1-DNA co-condensation drives DNA repair site assembly to prevent the disjunction of broken DNA ends ^30^. As the formation of chromatin-bound PARP1 is also triggered by PARPi such as talazoparib, it is possible that decitabine might have synergistic activity in dKRAS-PDAC with a broad spectrum of clinically available PARPi.

The rational application of these combination epigenetic approaches in clinic will also require a focus on predictive biomarker discovery to identify molecular and histologic determinants of susceptibility. A Phase I clinical trial of the DNA methyltransferase inhibitor decitabine in combination with the PARP inhibitor talazoparib has been conducted in relapsed/refractory acute myeloid leukemia ^31^, yet in unselected patients. The efficacy of the COMBO in dKRAS-/*BRCA1/2*^*wild-type*^ was here predicted using transcriptomic-based KRAS dependency scores as biomarkers of efficacy. We have previously demonstrated that assessing KRAS dependency in PDAC can be achieved using gene expression signature scores in clinical settings (NCT05360264) irrespective of specific KRAS mutations. Given the high prevalence of KRAS-dependent tumors ^4^, effective therapies against dKRAS-PDAC, such as COMBO, can offer enhanced therapies for a substantial proportion of PDAC patients.

Furthermore, we found that COMBO was significantly more effective than OLA alone when treating dKRAS-/*BRCA2*^*mut*^ PDAC cells (CAPAN-1), as demonstrated by two different combination index models showing strong synergistic antiproliferative activity across all analyzed concentrations and by the antimetastatic effect of COMBO in xenograft tumor models. These results suggest that PDAC tumors already candidates for OLA treatment, but classified as KRAS-dependent, might have higher therapeutic benefits from COMBO therapy than OLA treatment alone. There is also the question of whether KRAS dependency might be a molecular phenotype of other solid tumors, such as breast, ovary, and prostate cancer, which have a higher frequency of BRCA mutations than PDAC, and if DEC could also enhance the antitumor efficacy of OLA in these tumor subtypes.

We reported that common biomarkers in clinical use or under investigation for predicting response to PARP inhibitors (PARPi) do not fully predict the synergistic response to the combination of OLA and DEC compared to KRAS dependency status. Our findings, limited to the cellular models investigated, indicated that KRAS dependency was not associated with common homologous recombination deficiency (HRD) biomarkers or microsatellite instability in PDAC cell lines. Additionally, KRAS dependency scores were not associated with the RAD51 score. Therefore, KRAS dependency appears to be a novel, independent biomarker for predicting response to the OLA and DEC combination treatment in PDAC. Evidence suggests a molecular correlation between HR-repair pathways and KRAS dependency pathways in tumors, particularly in response to radiotherapy ^32^, and a correlation between KRAS and RAD51 has been observed in colon cancer ^33^. Further investigation into the molecular mechanisms underlying KRAS dependency and its interaction with DNA damage response pathways may provide valuable insights into patient stratification and treatment selection. Furthermore, based on the data presented, it is possible to speculate that combining KRAS dependency scores with biomarkers of HRD, such as genomic alterations of DNA repair genes or RAD51 score, may further refine the selection of patients most likely to respond to therapies with decitabine and PARPi, thus enhancing personalized therapy in PDAC.

## EXPERIMENTAL PROCEDURES

### Reagents and Chemicals

Refer to Supplementary Experimental procedures for details

### Cell Lines and Cell Authentication

CAPAN-1, BXPC3, ASPC1, and PaTu 8902 cell lines were provided by Dr. Giuseppe Diaferia (European Institute of Oncology, Milan, Italy). KP-4 and PaTu 8988t cell lines were provided by the cell culture facility at The University of Texas MD Anderson Cancer Center (UTMDACC; Houston, Texas). HPAF-II cell lines were provided by Dr. Michele Milella (Department of Medicine, University of Verona, Italy). The SU8686 cell line was purchased from ATCC. Human pancreatic ductal epithelia (HPDE) KRAS^*G12V*^ isogenic derivatives were obtained from Dr. D. Melisi (Department of Medicine, University of Verona, Italy). Cell line identities were genetically validated according to relative cell bank procedures and were cultured for fewer than 6 weeks after resuscitation. Additionally, full transcriptomic profiles from the cell lines under investigation were routinely analyzed and compared with reference transcriptomic profiles derived from multiple references in publicly available databases (GEO), to assess the global genetic identity. Upon passage 10, cells were discarded. All cells were routinely tested for Mycoplasma contamination using a MycoFluor Mycoplasma Detection Kit (Thermo Fisher Scientific).

PaTu 8902 and PaTu 8988t cells were maintained in DMEM high glucose containing L-glutamine and sodium pyruvate, supplemented with 10% FBS (Gibco). CAPAN-1 cells were grown in RPMI supplemented with 20% FBS (Gibco) plus 1% glutamine (Gibco). KP-4 cells were maintained in Iscove’s modified Dulbecco’s medium (Gibco), supplemented with 20% FBS and 200 mmol/L glutamine. SU8686, HPAF-II, BxPC-3, HPDE and ASPC1 cells were maintained in RPMI supplemented with 10% FBS (Gibco) plus 1% glutamine (Gibco). All culture growth media were supplemented with 10,000 U/mL penicillin–streptomycin (Gibco). HPDE-KRAS^*G12V*^ cells were grown in RPMI supplemented with 10% FBS (Gibco) plus 1% glutamine (Gibco).

### KRAS Dependency Signature Score Calculation Methods

The gene signature scores were calculated by subtracting the average normalized expression of “down” genes from the average normalized expression of “up” genes, defining the L-score and the S-score. The RNA sequencing data were normalized by taking the log10 ratio of expression of the sample compared to the average expression of the gene across all samples. The L-score genes are described in (8); the S-score genes are described in (9). The top KRAS-dependent genes were selected as “up” genes, and the top KRAS-independent genes were selected as “down” genes. The microarray data were normalized by performing the same transformation as described for the RNA sequencing data, except on the normalized intensity values rather than TPM. The L and S scores of dependency do not correlate with the cellular proliferation index as estimated by the correlation analysis of the L and S scores with cellular doubling time (Figure S2).

### Cell Viability Assay and Half-Maximal Inhibitory Concentration (IC50) Determination

The viability of cells was evaluated using the CellTiter-Glo Luminescent Cell Viability Assay (Promega). For KP-4, PaTu 8902, and PaTu 8988t, a total of 5 × 10^2 cells were seeded at T=0. For CAPAN-1, 1 × 10^3 cells were seeded onto 96-well plates. After 24 hours, cells were exposed to indicated serial concentrations of drugs, and culture media and treatments were refreshed after 72 hours. Viability was measured either 72 hours or 144 hours later, as indicated, using the CellTiter-Glo Luminescent Cell Viability Assay (Promega), according to the manufacturer’s instructions. Dose-response curves and half-maximal inhibitory concentration (IC50) values with 95% CI were generated using GraphPad Prism software (GraphPad Software).

### Comet assay

Refer to Supplementary Experimental procedures for details

### Cell cycle analysis by flow cytometry

Refer to Supplementary Experimental procedures for details

### Immunofluorescence analysis of DDR markers

Refer to Supplementary Experimental procedures for details

### RAD51 score

Cells (1500 cells/cm^2^) were plated on glass coverslips in 24-well plates for 96 hours. Then, cells were washed two times with 1X PBS and fixed using 4% formaldehyde for 20 minutes at room temperature. Fixed cells were washed two times with 1X PBS and permeabilized using 0.25% Triton X-100/1X PBS solution for 5 minutes. After permeabilization, cells were washed three times with 1X PBS and then blocked using 1% bovine serum albumin (Sigma)/1X PBS solution for 40 minutes at room temperature. Cells were washed three times with 1X PBS and then incubated with RAD51 primary antibody (ab133534, Abcam, Cambridge, UK) at a 1:500 dilution in PBS-BSA (1%) for 2 hours at room temperature. The cells were then washed three times with 1X PBS and incubated with geminin-L-CE primary antibody (Leica Biosystems) for 1 hour. Next, cells were washed three times with 1X PBS and incubated with secondary antibodies (α-mouse, two drops/ml in PBS-BSA1% and α-rabbit, two drops/ml in PBS-BSA1%) for 1 hour at room temperature. After three washes with 1X PBS, cells were mounted with Fluoroshield™ with DAPI (Sigma Aldrich) and coverslips were then left to dry in the dark. After 24 hours, stained cells were acquired using an Olympus AX70 microscope at 40x or 60x magnification.

For automated geminin-positive cell and nuclear foci quantification, a custom pipeline in CellProfiler was developed to measure and export raw data containing both the geminin-positive nuclei per image and the number of nuclear foci per geminin-positive cell. The RescaleIntensity and MedianFilter modules relatively change the intensity range of each image and reduce noise. Then, total nuclei and geminin-positive nuclei are identified as primary objects and related with the RelateObjects module to determine the ratio between total nuclei and geminin-positive nuclei and classify them into two different groups with the ClassifyObjects module. The enhancement of speckles with the EnhanceOrSuppressFeatures module allows for the identification of RAD51 foci in all the cells. Lastly, the RelateObjects module was used to count the number of foci per geminin-positive cell. Parameters such as pixel unit diameter, threshold smoothing scale, and threshold correction factor change depending on the cell line.

Geminin-positive cells with at least three foci were considered HR proficient, and the percentage of HR proficient cells versus the total number of geminin-positive cells was considered the RAD51 score. Statistical analyses were performed using GraphPad Prism 8 with a Kruskal-Wallis test within RAD51 scores of the RAD51-low group and the RAD51-high group and a 2-way ANOVA test between the two groups.

### Genomic Data Analysis and Microsatellite Stability Assessment

Genomic mutational profiles of the selected PDAC cell lines were analyzed using data from the COSMIC and cBioPortal databases. We specifically examined gene mutations and copy number alterations (CNAs) in genes associated with DNA damage response (DDR) and homologous recombination, which are known to influence OLA cytotoxicity. Microsatellite stability (MS) for each cell line was annotated using previous reports and analysis from the CCLE panel (Gandi et al.2019).

### Subcellular Protein Fractions

Refer to Supplementary Experimental procedures for details

### Total Cell Lysates

Refer to Supplementary Experimental procedures for details

### “Combenefit” Method for Drug Combination Index Analysis

The sensitivity was tested in a 7-day-long proliferation assay. Cells were seeded in 96-well culture plates in different numbers per well depending on the cell line to reach 80%-90% confluency of control wells at the end of the assay. The following day, serial dilutions of Decitabine (1:4 for Pa-Tu-8902 and HPAF-II, 1:2 for Capan-1), and Olaparib (1:3 for Pa-Tu-8902 and HPAF-II, 1:2 for Capan-1), were added to the cells in single treatment and combining each concentration of the two drugs in a 4×5 matrix, then the treatments were refreshed after 72h. Seven days after the first treatment, the cell viability was assessed by Cell Titer-Glo Luminescent Cell Viability (CTG) assay (Promega) and measured by Varioscan Lux plate reader. Viability measured for each treatment condition was normalized to untreated controls. Final data are an average of at least three biological replicates with similar results. After CTG analysis, synergistic/antagonistic/additive combinations were analyzed using Combenefit software, which provides synergy distribution plots by comparing experimental data to mathematical models (e.g., HSA model) of dose responses for additive/independent combinations. The HSA model assesses the efficacy of a drug combination by comparing it to the effect of the single most effective drug in the combination. It operates under the assumption that the effect of the drug combination should not exceed the effect of the most potent single drug. In brief, the software first reads each experimental dose response as a matrix of percentages of the control and each single-agent effect is fitted with a dose–response curve. If the observed effect of the drug combination is greater than the effect predicted by the HSA model, the combination is considered synergistic. If the observed effect matches the predicted effect, the combination is considered additive. If the observed effect is less than the predicted effect, the combination is considered antagonistic. Combenefit applies the HSA model to the input data. The software calculates the expected effect of the combination based on the highest single-agent effect. The results are typically presented in a graphical format (e.g., heatmaps). Combenefit may also provide statistical metrics to quantify the degree of interaction, helping users to make more precise interpretations of the data.

### “Calcusyn” Method for Drug Combination Index Analysis

Refer to Supplementary Experimental procedures for details

### Immunoblot Analysis

Refer to Supplementary Experimental procedures for details

### Xenograft Tumor Experiments

NOD.Cg-Prkdcscid Il2rgtm1Wjl/SzJ (NSG) mice were purchased from Jackson Laboratory and bred in the animal facility of CAST, G. D’Annunzio University, Chieti. Mice were housed on a 12-hour light/dark cycle and maintained at 24–26°C under pathogen-limiting conditions. Cages, bedding, and food were autoclaved before use, and food and water were provided ad libitum. Animal care and all experimental procedures were approved by the Ethics Committee for Animal Experimentation of the Institute according to Italian law (Authorization n°640/2022-PR). Seven-week-old mice were subcutaneously injected into the right flank with a suspension of 1.5×10^6 HPAF-II, KP-4, PaTu-8988t, and CAPAN-1 cells diluted in 200 μl of a 1:1 mixture of Matrigel (BD Biosciences) and PBS (Corning). Treatments were started when tumors reached palpable volumes. Decitabine, dissolved in saline, was administered intraperitoneally (IP) at 0.2 or 0.5 mg/kg body weight daily for the first week and three times a week for the following weeks. Olaparib, dissolved in 0.125% carboxymethylcellulose, was administered orally (OS) at 30 mg/kg body weight daily. Control mice received oral and IP treatments with vehicles only. Tumor volume was measured once or twice a week with calipers and calculated using the formula: V(mm^3) = L(mm) × W(mm)^2 / 2, where W is tumor width and L is tumor length. Mice were sacrificed when tumors in the control group reached 1 cm^3 in volume or were ulcerating, according to approved guidelines of the institution’s animal ethics committee. The health status of the animals was monitored daily, and body weight was measured at least once a week during treatments, with the percentage of body weight loss calculated.

### Histological and Immunohistochemical Analysis

Refer to Supplementary Experimental procedures for details

## Statistical Analysis

All in-vitro experiments were performed in triplicate wells and repeated multiple times using independent biological replicates (n). Student’s t-test was used to test for statistical significance of differences between groups and controls. P values < 0.05 were considered statistically significant. Sample correlations were calculated using Spearman’s correlation coefficient.

Statistical analysis for comet assays was performed using GraphPad Prism 8 and R. For comparison between two groups, a Mann-Whitney test was performed for the p-value and a Wilcoxon test was performed for the effect size. For comparisons among three or more groups, a Dunn test was performed for the p-value and a Wilcoxon test was performed for the effect size.

GraphPad Prism 8 was used for immunofluorescence nuclear foci quantification and group comparisons with an unpaired t-test. In the RAD51 score experiment, a Kruskal-Wallis test and two-way ANOVA were performed.

Statistical analyses for immunoblot analysis were performed using GraphPad Prism 8 and R with a two-way ANOVA test for comparisons between pairs of cell lines and one-way ANOVA with multiple comparisons through post-hoc Tukey HSD (Honestly Significant Difference) test within each experimental set of cell lines. In the graph, the same letter represents no statistical relevance in the difference between groups, while different letters represent a significant p-value (< 0.05) for the comparison between groups.

For xenograft-based model experiments, statistical differences between experimental groups were evaluated by applying a two-way ANOVA test or a two-tailed unpaired Student’s t-test. Differences were considered statistically significant when p-values were less than 0.05. All analyses were conducted using GraphPad Prism 5 software (GraphPad, San Diego, CA, USA).

## Supporting information

Supplementary experimental procedures

## Data Availability Statement

Genomic data for the pancreatic ductal adenocarcinoma (PDAC) cell lines used in this study are publicly accessible through the Cancer Cell Line Encyclopedia (CCLE) at the CCLE Portal. Additional mutation and copy number alteration data were obtained from the COSMIC (Catalogue Of Somatic Mutations In Cancer) database, accessible at https://cancer.sanger.ac.uk/cosmic, and the cBioPortal for Cancer Genomics, accessible at https://www.cbioportal.org/. Any other data supporting the findings of this study are available from the corresponding author upon request.

## SUPPLEMENTAL INFORMATION

Supplemental information includes Supplemental Experimental Procedures and ten figures and one table with legends:

Figures S1–S10

Legends to supplementary Figures

Table S1

Legend to Table S1

Supplemental Experimental Procedures

## ACKNOWLEDGMENTS

We thank Dr. Giuseppe Diaferia (IFOM, Milan, Italy) for providing some PDAC cell lines. We also thank Dr. A. Cavazzani (University of Tor Vergata, Rome) and Dr. C. Angelini (IAC-CNR) for their support in statistical analysis. We are grateful to Dr. V. Marabitti for technical advice on comet assay protocols, and Dr. A. Torcinaro (IBBC-CNR) for advice on CellProfiler software. We thank Dr. C. di Pietro (IBBC-CNR) for technical support with confocal microscopy, and Dr. G. Pfister (EMBL-Monterotondo, Rome) and Dr. A. Giovinazzo (IBBC-CNR) for advice and support with flow cytometry analysis.

G.A. and M.F. were supported by fellowships from Associazione “Nastro Viola”, Italy. This work was supported by LILT, Intramural grants of IRE, and Associazione “Nastro Viola”.

## AUTHOR CONTRIBUTIONS

G.A., M.F., and A.L. conducted experiments, developed methodologies, and analyzed data, including statistical analysis, as well as interpreted results. M.F. specifically performed imaging analysis, while C.M. and F.dM. assisted in certain experiments. M.M and M.F. performed the computational analysis of KRAS-dependency scores. M.I. oversaw in vivo studies, contributed to discussions, data analysis, and methodology development. L.C., G.A., and M.F designed and conceived the study. L.C. supervised the study, edited the manuscript draft, and provided overall guidance and financial support for the project. All authors thoroughly reviewed and contributed to the final version of the manuscript.

