## Supplementary material for "Enhancing PDAC Therapy: Decitabine-Olaparib Synergy Targets KRAS-Dependent Tumors": Legends to supplementary figures and table

**Figure S1.** Correlation plot comparing the Sscores versus Lscores of KRAS-dependency in 41 pancreatic cell lines calculated from the RNA sequencing data released by the CCLE Consortium**.**

**Figure S2 (A-B).** Correlation plot comparing the Lscores (A) or the Sscores of KRAS-dependency versus the doubling time in 41 pancreatic cell lines calculated according to data released by the CCLE Consortium or cell line provider information (https://www.cellosaurus.org).

**Figure S3.** Representative images of immunofluorescence from experiments in Figure 1A. Immunofluorescence analysis was performed to assess the activation of DNA damage response (DDR) markers in dKRAS- (BxPC-3) and indepKRAS- (KP-4) PDAC cell lines following treatment with decitabine (DEC) or vehicle (DMSO) for 72 hours. Cells were stained for phosphorylated H2AX (γH2AX), XRCC1, RAD51, and 53BP1 markers. The nuclei were counterstained with DAPI. Scale bars, 10 μm for magnification x60. Scale bars, 20 μm for magnification x40. The inset represents a further magnification of a single cell image.

**Figure S4. ATR or ATM inhibitors suppress DEC-induced phosphorylation of CHK1 at Ser345 and CHK2 at Thr68 in dKRAS-PDAC cells (PaTu 8902 and HPAF-II).** Histograms represent the densitometric analysis of γCHK1/CHK1 (A) and γCHK2/CHK2 (B) from immunoblot experiments described in Figure 1G. Quantification of immunoblots of total proteins extracted from the indicated cell lines treated with DMSO or DEC (100 nM) for 72 hours. Where indicated, the ATM inhibitor (ATMi) AZD1390 (10 nM) or the ATR inhibitor (ATRi) BAY-1895344 (2 µM) was added 1 hour before harvesting the cells. Histograms represent mean ± SD (error bars) from triplicate experiments. Statistical analyses were performed using GraphPad Prism 8 and R with one-way ANOVA with multiple comparisons through post-hoc Tukey HSD (Honestly Significant Difference) test within the experimental set of each cell line (same letter in the graph represents no statistical significance in the difference between groups; different letters represent a significant p-value (< 0.05) for the comparison between groups).

**Figure S5.** (A) Activation of DDR markers was significantly enhanced by combined DEC and OLA treatment in dKRAS cells, but not in indepKRAS PDAC cells. Histograms represent the densitometric analysis of γCHK1/ CHK1 (upper panels) and γH2AX/actin (lower panel) from immunoblot experiments described in Figure 2D. (B) Densitometric analysis of total PAR immunoblot of the nuclear soluble (NS) protein fraction experiments, as described in Figure 2E (left panels). RCC1 protein was used as a loading control. (C-E) Histograms represent the densitometric analysis of immunoblot experiments performed as described in Figure 2E (right panels). Histograms represent PARP1 (C) and XRCC1 (D) protein quantification over H3 immunoblot used as a loading control for the nuclear insoluble (NI) protein fraction (upper panels) or over RCC1 immunoblot used as a loading control for the nuclear soluble (NS) fraction (lower panels). Quantification of γCHK1/CHK1 is also presented (E). All histograms represent mean ± SD (error bars) from triplicate experiments. Statistical analyses were performed using GraphPad Prism 8 and R with a one-way ANOVA with multiple comparisons through post-hoc Tukey HSD (Honestly Significant Difference) test within the experimental set of each cell line (same letter in the graph represents no statistical significance in the difference between groups; different letters represent a significant p-value (< 0.05) for the comparison between groups).

**Figure S6. Determination of DEC and OLA half-maximal inhibitory concentration (IC_50_)** **in selected PDAC cell lines.** The indicated cell lines were plated and grown for 6 days with different doses of decitabine, ranging from 0.000025 μmol/L to 1000 μmol/L, or 0.0005 μmol/L to 180 μmol/L of OLA. Cell viability was then assayed by the ATP-based CellTiter-Glo assay. Graphs report the percentage of inhibition compared with DMSO (vehicle)-treated cells (n = 5).

**Figure S7.** **Combination index (CI) for the interaction between DEC and OLA obtained with the Calcusyn model.**

Upper panels: Combination index (CI) values plotted as a function of the fraction affected (Fa). The CI values of < 1 (below the lower dashed line), 1-1.1, and > 1.1 (above the upper dashed line) represent synergism, additivity, and antagonism, respectively. Circles indicate experimental points. Lower panels: Summary of the activity fraction affected (Fa) ranging from 0.5 to 0.95, CI values, and doses of DEC, OLA, and COMBO as analyzed by Calcusyn software. Data are representative of four to five independent experiments with equivalent results. Concentrations tested ranged from 0.25x to 4x of the IC_50_ concentrations and were used at a constant ratio. An r value of 0.95 or above indicated good conformity of the dose-effect data with respect to the median-effect principle. The r values for all experiments were 0.99 or higher. Drug combination analysis showed that OLA and DEC were highly synergic to additive in dKRAS-/*BRCA1/2^wild-type^* PDAC cell lines within the tested concentrations, with a trend of antagonism (CI > 1) when both drugs were used at the highest concentration. Of note, DEC and OLA were highly synergic in dKRAS-/*BRCA2^mut^ PDAC* cells (CAPAN-1) within all the tested drug concentrations.

**Figure S8.** Quantification of comet tail moment (TM) was performed in dKRAS-PDAC cells (PaTu 8902, HPAF-II, and CAPAN1) and indepKRAS-PDAC cells (PaTu 8988t and KP-4) following 72-hour treatment with DMSO, low-dose DEC, OLA, or simultaneous DEC plus OLA (COMBO) at the indicated drug concentrations selected based on Combination Index results (Figure 3A). Data are presented as the median from a representative replicate experiment of the alkaline comet assay. At least 200 nuclei were counted for each replicate. To quantify the magnitude of treatments-induced DNA damage among groups, the effect size (e.s.) was estimated by the Wilcoxon test by three biological replicates. The Rstatix package was implemented for the calculation of the effect size. Statistical significance among the indicated comparison groups was calculated using the Dunn’s test. Statistical significance: ****, p < 0.0001. Medium (M) to large (L) effect sizes suggest substantial DNA damage, while small (S) effect sizes indicate minimal to no response to treatment. Effect size (e.s): S = Small; M = Medium; L = Large. The calculated effect sizes indicate a robust induction of DNA damage by COMBO compared to single drugs in dKRAS- cells. No significant increase in TM was observed in indepKRAS cells following COMBO treatment with effect sizes calculated as small (S). Results of the neutral comet assay are reported in Figure 3B.

**Figure S9.** DDR markers were activated by COMBO treatment, compared to low-dose DEC or OLA alone, in dKRAS-/BRCA1/2 wild-type PDAC cell lines (PaTu 8902, HPAF-II) and dKRAS-/BRCA2 mutant PDAC cells (CAPAN-1) but not in indepKRAS-/BRCA1/2 wild-type PDAC cell lines (PaTu 8988t and KP-4). (A-D) Densitometry analysis of immunoblot experiments as reported in Figure 3D. The indicated cell lines were treated for 72 hours with DMSO, DEC, OLA, or COMBO at drug concentrations as in Figure 3B. Histograms report protein levels normalized as follows: total PAR/actin (A), γCHK1(Ser345)/CHK1 (B), γRPA32(Ser4/Ser8)/actin (C), and γH2AX(Ser139)/actin (D). All histograms represent mean ± SD (error bars) from triplicate experiments. Statistical analyses were performed using GraphPad Prism 8 and R with a one-way ANOVA with multiple comparisons through post-hoc Tukey HSD (Honestly Significant Difference) test within the experimental set of each cell line (same letter in the graph represents no statistical significance in the difference between groups; different letters represent a significant p-value (< 0.05) for the comparison between groups).

**Figure S10.** (A) Mice survival expressed as a percentage on Days 7, 9, and 28 after treatments with different doses of Decitabine (0.5mg/kg or 0.2 mg/kg), Olaparib (30mg/kg), and their combinations. (B-E) Tumor growth kinetics of each mouse in each treatment group for HPAF-II cell line (B), KP-4 cell line (C), PaTu 8988t cell line (D), and CAPAN-1 cell line (E). (F-I) Percentage of change in body weight during the treatment of mice injected with HPAF-II cells (F), KP-4 cells (G), PaTu 8988t cells (H), and CAPAN-1 cells (I).

**LEGEND TO SUPPLEMENTARY TABLE**

**TABLE S1**. List of S-scores and L-scores of KRAS-dependency of 41 pancreatic cell lines calculated from the RNA sequencing data released by the CCLE Consortium and used to generate Figure S1. Cell lines are ranked according to decreasing LScore value.
