## Supplementary experimental procedures for "Enhancing PDAC Therapy: Decitabine-Olaparib Synergy Targets KRAS-Dependent Tumors"

**Reagents and Chemicals**

Decitabine (DEC), 5-Aza-2’-deoxycytidine (A3656), Dimethyl Sulfoxide (DMSO), Protease Inhibitor Cocktail, Bovine Serum Albumin, and β-Nicotinamide adenine dinucleotide (NAD^+^) No. N8410 were purchased from Sigma-Aldrich (St. Louis, MO, USA). Olaparib (OLA): AZD2281, Ku-0059436 (S1060), AZD1390, ATM Inhibitor (ATMi), and BAY-1895344, ATR Inhibitor (ATRi), were purchased from Selleckchem. Poly(ADP-ribose) Glycohydrolase (PARG) inhibitor: PDD00017273, was purchased from MedChemExpress. FxCycle™ PI/RNase Staining Solution was purchased from Thermo Fisher Scientific (Waltham, MA, USA). CellTiter-Glo Assay Reagent was purchased from Promega.Fetal Bovine Serum (FBS), Penicillin/Streptomycin (P/S), and 1% Glutamine were purchased from Gibco.

Antibodies are listed in each specific experimental section.

**Comet assay**

Cell Preparation

Cells (3000 cells/cm²) were plated in 100mm dishes and treated after 24 hours as indicated. Cells were either used immediately or frozen in freezing medium in Eppendorf tubes and stored at -80°C. When needed, cells were thawed at 37°C, centrifuged at 4°C for 10 minutes (1500 rpm), washed in 1x PBS, and centrifuged again. The cell suspension in 1x PBS (10 µl/50k cells) was combined with 75 µl of 0.5% low melting point agarose (LMPA) at 37°C, gently mixed, and rapidly spread (75 µl) onto a pre-coated glass slide with normal melting point agarose (NMPA). Slides were immediately covered with coverslips and the agarose was allowed to solidify at 4°C for about 5 minutes.

**Alkaline Comet Assay.** Coverslips were gently removed, and slides were immersed in alkaline lysis solution for at least 18-20 hours (no more than 5 days) at 4°C in the dark. Slides were then transferred into a horizontal electrophoresis tank containing freshly prepared electrophoresis buffer for 20 minutes at 4°C to allow DNA to unwind. Electrophoresis was performed at 0.6 V/cm (cm refers to the diagonal between the electrodes) and 250-300 mA for 20 minutes in the same alkaline solution. Slides were rinsed with Neutralization Buffer three times for 5 minutes each, washed with MilliQ-H₂O, fixed with ice-cold 100% ethanol, and dried at room temperature in the dark.

**Neutral Comet Assay.** Coverslips were gently removed, and slides were immersed in freshly prepared neutral lysis solution for 45 minutes at 4°C in the dark. Slides were washed in 1x TBE at 4°C and transferred into a horizontal electrophoresis tank containing 1x TBE at 4°C. Electrophoresis was performed at 0.6 V/cm (cm refer to the diagonal between the electrodes) and 6-7 mA for 15 minutes. Slides were rinsed with MilliQ-H₂O, fixed with ice-cold 100% ethanol, and dried at room temperature in the dark.

Slides were stained with Syber Safe diluted in MilliQ-H₂O, and after at least 12 hours, observed for image acquisition using an Olympus AX70 microscope at 20x magnification. Images of at least 200 cells per sample were captured for analysis. Comet images were analyzed using Comet Score software to measure tail moment.

Statistical analyses were performed with GraphPad Prism 8 and R. For comparison between two groups, a Mann-Whitney test was performed for the p-value, and a Wilcoxon test was performed for the effect size. For comparisons between three or more groups, a Dunn test was performed for the p-value, and a Wilcoxon test was performed for the effect size.

**Solutions**

- Alkaline Lysis Solution: 2.5 M NaCl, 100 mM EDTA, 10 mM Tris base, 4g NaOH (pH 10) in MilliQ-H₂O. 1 hour before sample preparation, add 1% Triton X-100 and keep at 4°C.
- Electrophoresis Buffer: 30 ml 10N NaOH, 5 ml 200 mM autoclaved EDTA (pH 10), complete to 1L with dH₂O and keep at 4°C.
- Neutralization Buffer: 0.4 M Tris-HCl (pH 7.5) in dH₂O, autoclave, and keep at 4°C.
- Neutral Lysis Solution: 6 ml 0.5 M EDTA (pH 8), 5 ml 10% SDS, complete to 100 ml with MilliQ-H₂O.
- TBE 10x: 108g Tris-base, 55g boric acid in 800 ml dH₂O, add 40 ml 0.5 M EDTA (pH 8), adjust pH to 8.5 and complete to 1L, autoclave, and store at room temperature.

**Cell cycle analysis by flow cytometry**

Cells were seeded in 100mm cell culture dishes. Twenty-four hours after plating, PaTu 8902 and PaTu 8988t were treated with DMSO (vehicle), 0.025 µM 5-aza-2'-deoxycytidine (Decitabine), 1.37 µM Olaparib, or a combination of both drugs for 72 hours. HPAF-II cells were treated with DMSO (vehicle), 0.045 µM Decitabine, 1.4 µM Olaparib, or a combination of both drugs for 72 hours. CAPAN-1 and KP4 cells were treated with DMSO (vehicle), 0.09 µM Decitabine, 0.5 µM Olaparib, or a combination of both drugs for 72 hours. Approximately 1 × 10^6 cells were harvested, washed in 1x PBS, fixed with cold 70% ethanol, and kept at -20°C for 24 hours. Fixed cells were re-suspended in FxCycle™ PI/RNase Staining Solution (Thermo Fisher Scientific) for 30 minutes before analysis by flow cytometry. Samples were acquired using either a FACScalibur Flow Cytometer (Becton Dickinson Company) or an Attune NxT Acoustic Focusing Cytometer (Thermo Fisher Scientific). The fcs files were analyzed using FlowJo software.

### Immunofluorescence analysis of DDR markers

### Cells (3000 cells/cm²) were plated on glass coverslips in 24-well plates and treated after 24 hours with 1.25 µM DEC for the indicated time. Cells were washed twice with 1x PBS and fixed with 4% formaldehyde for 20 minutes at room temperature. Fixed cells were washed twice with 1x PBS and permeabilized using 0.25% Triton X-100/1x PBS solution for 5 minutes. After permeabilization, cells were washed three times with 1x PBS and then blocked with 1% bovine serum albumin (Sigma)/1x PBS solution for 40 minutes at room temperature. Cells were washed three times with 1x PBS and then incubated with primary antibody. After washing three times with 1x PBS, cells were incubated with secondary antibodies. After three more washes with 1x PBS, cells were mounted with Fluoroshield™ with DAPI (Sigma-Aldrich) and allowed to dry in the dark. After 24 hours, stained cells were acquired using an Olympus AX70 microscope at 40x or 60x magnification. For quantitative analysis, images were processed using CellProfiler, and nuclear foci were quantified in relation to the total number of cells. A custom pipeline in CellProfiler was developed to measure and export raw data containing both the total nuclei and foci per image, and the number of nuclear foci per cell. Parameters such as pixel unit diameter, threshold smoothing scale, and threshold correction factor were adjusted depending on the cell line. The number of foci per nucleus in treated cells was compared to that in control cells, with at least 50 cells per group used for statistical analysis with an unpaired t-test using GraphPad Prism 8 software. For confocal microscopy, primary antibodies were incubated one at a time, while secondary antibodies were incubated simultaneously. Imaging analysis was performed using a Leica SP5 confocal microscope.

### Primary Antibodies Used in Immunofluorescence Analysis:

### Phospho-Histone H2A.X (Ser139) (Cat#05-636, Merck Millipore)

### XRCC1 Antibody (Cat#2735, Cell Signaling Technology)

### Anti-Rad51 antibody (ab133534, Abcam)

### 53BP1 Antibody (NB100-305, Novus Biologicals)

### Phospho-Chk1 (Ser345) (133D3), (Cat#2348, Cell Signaling Technology)

### PARP (46D11) (Cat#9532, Cell Signaling Technology)

### Secondary Antibodies Used in Immunofluorescence Analysis:

### Alexa Fluor 488 donkey anti-rabbit antibody (R37118, Thermo Fisher Scientific)

### Alexa Fluor 594 donkey anti-mouse antibody (R37115, Thermo Fisher Scientific)

**Subcellular Protein Fractions**

Cells were washed in cold phosphate-buffered saline 1X (PBS1X) and then collected by scraping in phosphate-buffered saline 1X (PBS1X) containing 1X PIC (Protease inhibitor cocktail, Cat#P8340, Sigma-Aldrich, St. Louis, MO, USA). Cells were collected in Eppendorf tubes and centrifuged at 500xg at 4°C, the supernatant was removed, and the Nuclear Isolation Buffer (NIB) [15mM Tris-HCl pH 7.5, 60mM KCl, 15mM NaCl, 5mM MgCl2, 1mM CaCl2, 250mM Sucrose, 1mM DTT, 2mM NaV, 1X Protease Inhibitor Cocktail, 1X PMSF, 1mM PARGi, 0.1% NP-40] was added to the cell pellet and incubated on ice for 10 minutes. The resuspended pellets were centrifuged at 2000xg at 4°C for 5 minutes and the supernatant (Cytoplasmic fraction) was collected in a clean tube and frozen in dry ice. The nuclei were gently washed twice with NIB buffer without detergent (NP-40) and then centrifuged at 2000xg for 5 minutes at 4°C. Nuclear Lysis Buffer (NLB) [20mM Tris-HCl pH 7.9, 300mM NaCl, 1.5mM MgCl2, 0.2mM EDTA, 0.5mM DTT, 1mM NaV, 1X Protease Inhibitor Cocktail, 1X PMSF, 1mM PARGi, 10% glycerol] was added to the pellet (nuclei) and incubated on ice for 30 minutes. After incubation, the supernatant (Nuclear Soluble fraction) was transferred to a clean tube and frozen in dry ice. NLB buffer complemented with 3µl/100µl of Micrococcal Nuclease and 5µl/100µl of CaCl2 were added to the pellet, vortexed at maximum power for 15 seconds and incubated at 37°C for 5 minutes, then vortexed again and centrifuged at 16000xg for 10 minutes at 4°C. The obtained pellet was lysed with radioimmunoprecipitation assay buffer 1X (RIPA-1: 50 mM Tris-HCl [pH 7.8], 150 mM NaCl, 5 mM EDTA, 15 mM MgCl2, 1% Nonidet P-40, 0.5% sodium deoxycholate, 1 mM dithiothreitol, 1X protease inhibitors, 1X PMSF, 50mM NaF, 10 mM β-glycerophosphate and 1 mM Na3VO4), incubated on ice for 10 minutes, centrifuged at 14000xg for 10 minutes at 4°C, and then the supernatant (Nuclear Insoluble) was transferred to a clean tube and frozen in dry ice. The cleared protein extracts of every cell fraction were quantified using the Bradford method (Bio-Rad).

**Total Cell Lysates**

Cells were washed in phosphate-buffered saline 1X (PBS1X) containing 1X protease inhibitor cocktail (Protease inhibitor cocktail, Cat#P8340, Sigma-Aldrich, St. Louis, MO, USA) and were lysed with a cell scraper in modified radioimmunoprecipitation assay buffer (RIPA-1: 50 mM Tris-HCl [pH 7.8], 150 mM NaCl, 5 mM EDTA, 15 mM MgCl2, 1% Nonidet P-40, 0.5% sodium deoxycholate, 1 mM dithiothreitol, 1X protease inhibitors, 1X PMSF, 50mM NaF, 10 mM β-glycerophosphate and 1 mM Na3VO4), and then immediately frozen in dry ice. Cleared protein extracts were obtained after sonication with Vibra-Cell™ Ultrasonic Liquid Processor VCX 500/VCX 750 (Sonics & Materials, Inc., Newtown, CT, USA) followed by centrifugation at 14000xg for 10 minutes at 4°C, and then the cleared protein extracts were quantified using the Bradford method (Bio-Rad).

**"Calcusyn" Method for Drug Combination Index Analysis**

Cells were treated with a combination of Ola and DEC using the method of constant ratio drug combination. The two drugs were used at a constant ratio of their concentrations. The concentrations used corresponded to 0.25, 0.5, 1, 2, and 4 times the IC50 of each agent (Ola: IC50=5.5µM; DEC: IC50=100nM for Pa-Tu-8902, Ola: 7.8µM; DEC: IC50=180nM for HPAF-II, Ola: 1µM; DEC: IC50=90nM for Capan-1). This method, using the combination index (CI) equation, allows quantitative determinations of drug interactions at increasing levels of cell kill. The CI value allows classification of the anti-tumor activity of the drug combination: a CI of less than, equal to, or more than 1 indicates synergistic, additive, or antagonistic effects, respectively. Fa is the fraction of cell death induced by drug treatment and ranges from 0 to 1, with 0 meaning no cell killing and 1 representing 100% cell killing. The sensitivity was tested in a 7-day-long proliferation assay. Cells were seeded in 96-well culture plates in different numbers per well depending on the cell line to reach 80%-90% confluency of control wells at the end of the assay. After 24 hours, cells were treated with serial dilutions (1:2) of each drug alone or in combination at a constant ratio (Ola) in three independent experiments with triplicate samples, and the treatments were refreshed after 72h. Seven days after the first treatment, the cell viability was assessed by Cell Titer-Glo Luminescent Cell Viability (CTG) assay (Promega) and measured by Varioscan Lux plate reader. Viability measured for each treatment condition was normalized to untreated controls. Final data are an average of at least three biological replicates with similar results. To determine the nature (synergism, additivity, and antagonism) of Ola and DEC interaction, we used the method proposed by Chou and Talalay using the Compusyn software**.**

**"Combenefit" Method for Drug Combination Index Analysis**

The sensitivity was tested in a 7-day-long proliferation assay. Cells were seeded in 96-well culture plates in different numbers per well depending on the cell line to reach 80%-90% confluency of control wells at the end of the assay. The following day, serial dilutions of Decitabine (1:4 for Pa-Tu-8902 and HPAF-II, 1:2 for Capan-1), and Olaparib (1:3 for Pa-Tu-8902 and HPAF-II, 1:2 for Capan-1), were added to the cells in single treatment and combining each concentration of the two drugs in a 4x5 matrix, then the treatments were refreshed after 72h. Seven days after the first treatment, the cell viability was assessed by Cell Titer-Glo Luminescent Cell Viability (CTG) assay (Promega) and measured by Varioscan Lux plate reader. Viability measured for each treatment condition was normalized to untreated controls. Final data are an average of at least three biological replicates with similar results. After CTG analysis, synergistic/antagonistic/additive combinations were analyzed using Combenefit software, which provides synergy distribution plots by comparing experimental data to mathematical models (e.g., HSA model) of dose responses for additive/independent combinations. The HSA model assesses the efficacy of a drug combination by comparing it to the effect of the single most effective drug in the combination. It operates under the assumption that the effect of the drug combination should not exceed the effect of the most potent single drug. In brief, the software first reads each experimental dose response as a matrix of percentages of the control and each single-agent effect is fitted with a dose–response curve. If the observed effect of the drug combination is greater than the effect predicted by the HSA model, the combination is considered synergistic. If the observed effect matches the predicted effect, the combination is considered additive. If the observed effect is less than the predicted effect, the combination is considered antagonistic. Combenefit applies the HSA model to the input data. The software calculates the expected effect of the combination based on the highest single-agent effect. The results are typically presented in a graphical format (e.g., heatmaps). Combenefit may also provide statistical metrics to quantify the degree of interaction, helping users to make more precise interpretations of the data.

**Immunoblot Analysis**

Protein samples were separated on 4%–12% or 4%-20% Tris-Glycine gels (Novex™ WedgeWell™, Invitrogen, Waltham, MA, USA) and transferred to nitrocellulose membranes (Amersham™ Protran™ 0.45µm NC). Membranes were blocked in TBS-T (0.1% Tween-20) containing 5% bovine serum albumin, incubated with primary antibodies according to the manufacturer's instructions, and then incubated with horseradish peroxidase-conjugated goat anti-rabbit IgG (Cat#7074, Cell Signaling Technology, Inc., Danvers, MA, USA) or anti-mouse IgG (Bethyl Laboratories, A90-116P). Detection was performed using enhanced chemiluminescence (ECL SuperSignal™ West Femto Maximum Sensitivity Substrate, Thermo Scientific, Waltham, MA, USA). The following primary antibodies were used for Western blotting: PARP1 (46D11) Rabbit mAb (Cat#9532, Cell Signaling Technology, Inc.), PAR/pADPr Antibody (Cat#4335-MC-100, R&D Systems, Minneapolis, MN, USA), XRCC1 Antibody (Cat#2735, Cell Signaling Technology, Inc.), β-Actin antibody (No. SAB5600204, Sigma-Aldrich, St. Louis, MO, USA), Histone H3 (D1H2) XPR Rabbit mAb (Cat#4499, Cell Signaling Technology, Inc.), CHK1 (2G1D5) Mouse mAb (Cat#2360, Cell Signaling Technology, Inc.), phospho-RPA32 (S4/S8; Bethyl Laboratories, A300-245A), phospho-Chk1 (Ser345) (133D3) Rabbit mAb (Cat#2348, Cell Signaling Technology, Inc.), phospho-Chk1 (S317) (Cat#2344, Cell Signaling Technology, Inc.), phospho-Chk2 (Thr68) (Cat#2661, Cell Signaling Technology, Inc.), Chk2 (1C12) Mouse mAb (Cat#3440, Cell Signaling Technology, Inc.), phospho-Histone H2AX (Ser139) (Cat#05-636, Merck Millipore, Burlington, MA, USA), and RCC1 (N-19) (sc-1161, Santa Cruz Biotechnology, Inc., Santa Cruz, CA, USA). Statistical analyses were performed using GraphPad Prism 8 with a two-way ANOVA test for comparisons between pairs of cell lines, and one-way ANOVA with multiple comparisons through post-hoc Tukey HSD (Honestly Significant Difference) Test within each experimental set of cell lines. In the graph, the same letter represents no statistical relevance in the difference between groups, while different letters represent a p-value < 0.05 for the difference between groups.

**Histological and Immunohistochemical Analysis**

Primary tumors and lungs were fixed in 10% neutral buffered formalin, embedded in paraffin, and sectioned at a 5-µm thickness for histological analysis. To optimize the detection of microscopic metastases and ensure systematic uniform and random sampling, lungs were cut transversally into 2.0-mm thick parallel slabs, with a random position of the first cut within the first 2 mm of the lung, resulting in 5-8 slabs per lung. The slabs were then embedded cut surface down and sections were stained with Hematoxylin and Eosin. Slides were independently evaluated by two pathologists to quantify the number of tumor lesions. Immunohistochemical staining on formalin-fixed paraffin-embedded (FFPE) tissue from tumor xenografts was performed using the following primary antibody: mouse monoclonal anti-pH2AX antibody (1/500 dilution; 05-636, Millipore). Antigen retrieval was performed by microwaving slides for 10 minutes in Tris-EDTA buffer (pH 9.0, Sigma Aldrich). After blocking for 10 minutes with protein block (Dako) at room temperature, sections were incubated for 30 minutes with anti-pH2AX. Envision anti-mouse IgG (Dako) was used as the secondary antibody for 30 minutes, followed by 3,3'-diaminobenzidine (DAB) as the chromogenic agent. After chromogen incubation, slides were counterstained in Hematoxylin (Bio-Optica) and images were scanned with a Nanozoomer scanner from Hamamatsu. Quantification of pH2AX (% of positive cells) was performed on whole tumor sections (5-6 samples per group) analyzed with QuPath 0.3.2 software using the positive cell detection tool. The percentage of positive cells was plotted using GraphPad Prism 5 software**.**
