## Supplementary figures 1-10 for "Enhancing PDAC Therapy: Decitabine-Olaparib Synergy Targets KRAS-Dependent Tumors"

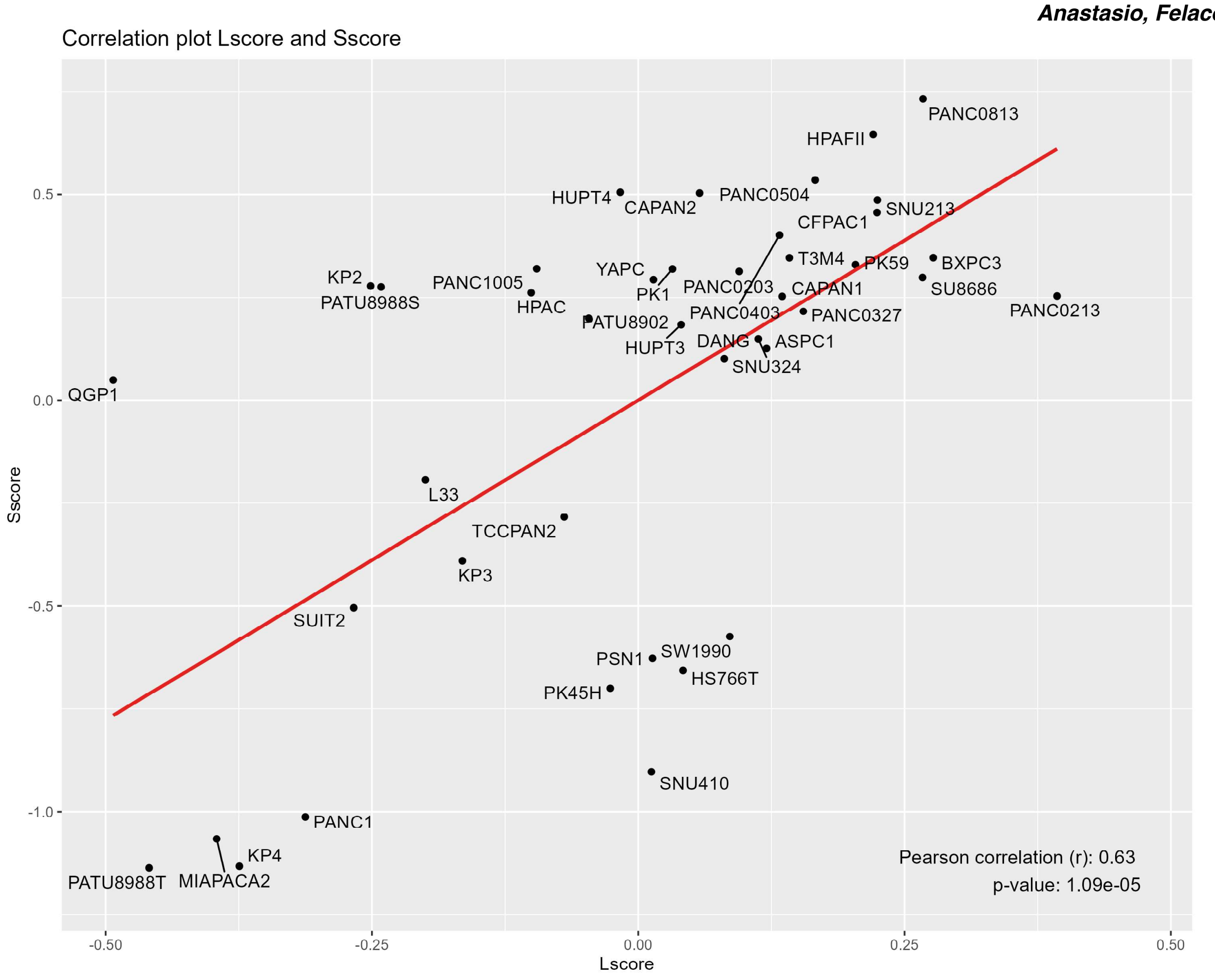
Correlation plot Lscore and Sscore
Sscore
Lscore
0.5
0.0
-0.5
-1.0
-0.50
-0.25
0.00
0.25
0.50
PANC0813
HPAFII
HUPT4
CAPAN2
PANC0504
SNU213
CFPAC1
T3M4
PK59
BXPC3
SU8686
PANC0213
KP2
PATU8988S
PANC1005
HPAC
YAPC
PK1
PANC0203
CAPAN1
PANC0403
PANC0327
PATU8902
HUPT3
DANG
ASPC1
SNU324
QGP1
L33
TCCPAN2
KP3
SUIT2
SW1990
PSN1
HS766T
PK45H
SNU410
PANC1
KP4
PATU8988T
MIAPACA2
Pearson correlation (r): 0.63
p-value: 1.09e-05

### Figure S1

# A

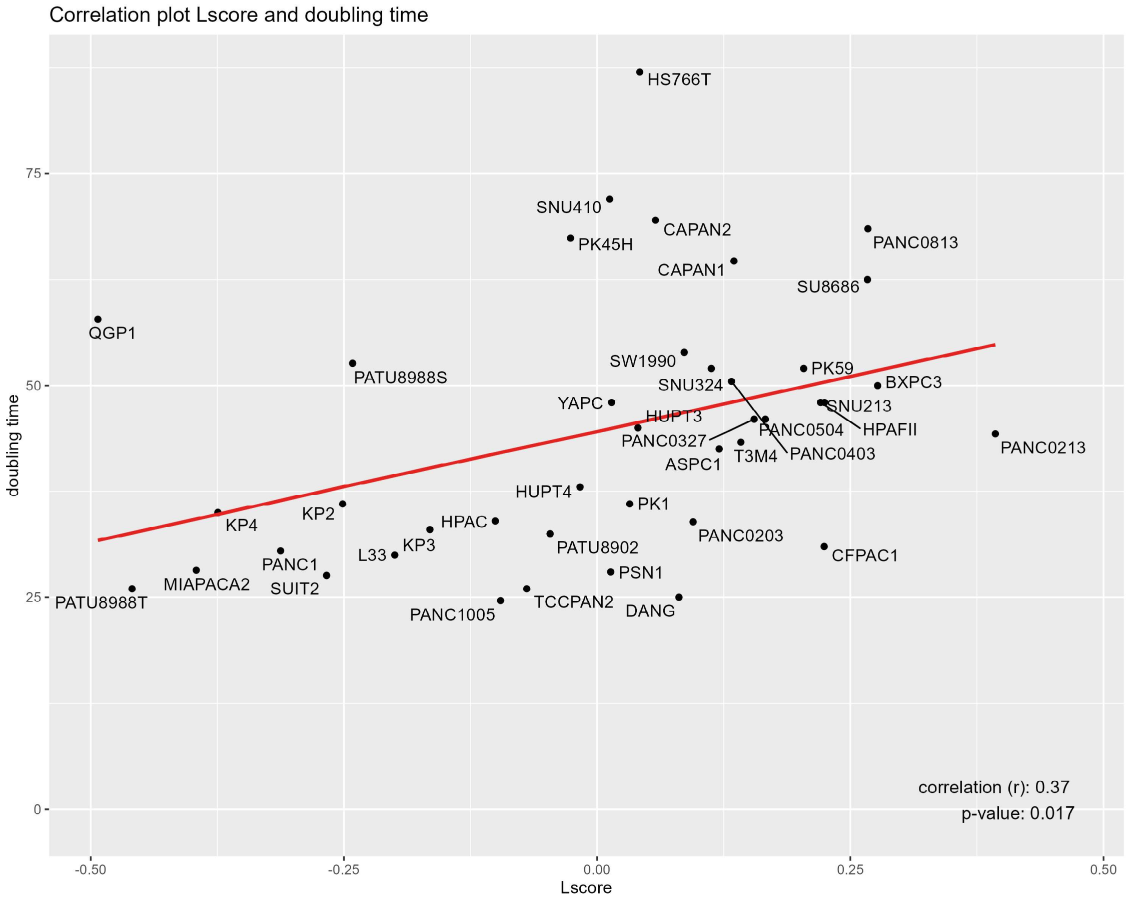

Correlation plot Lscore and doubling time
doubling time
Lscore
0
25
50
75
-0.50
-0.25
0.00
0.25
0.50
HS766T
SNU410
CAPAN2
PK45H
PANC0813
CAPAN1
SU8686
QGP1
SW1990
PK59
PATU8988S
SNU324
BXPC3
YAPC
SNU213
HUPT3
PANC0504
HPAFII
PANC0327
T3M4
PANC0403
PANC0213
ASPC1
HUPT4
PK1
KP2
KP4
HPAC
PANC0203
KP3
PATU8902
L33
CFPAC1
PANC1
PSN1
MIAPACA2
SUIT2
PATU8988T
TCCPAN2
DANG
PANC1005
correlation (r): 0.37
p-value: 0.017


# B

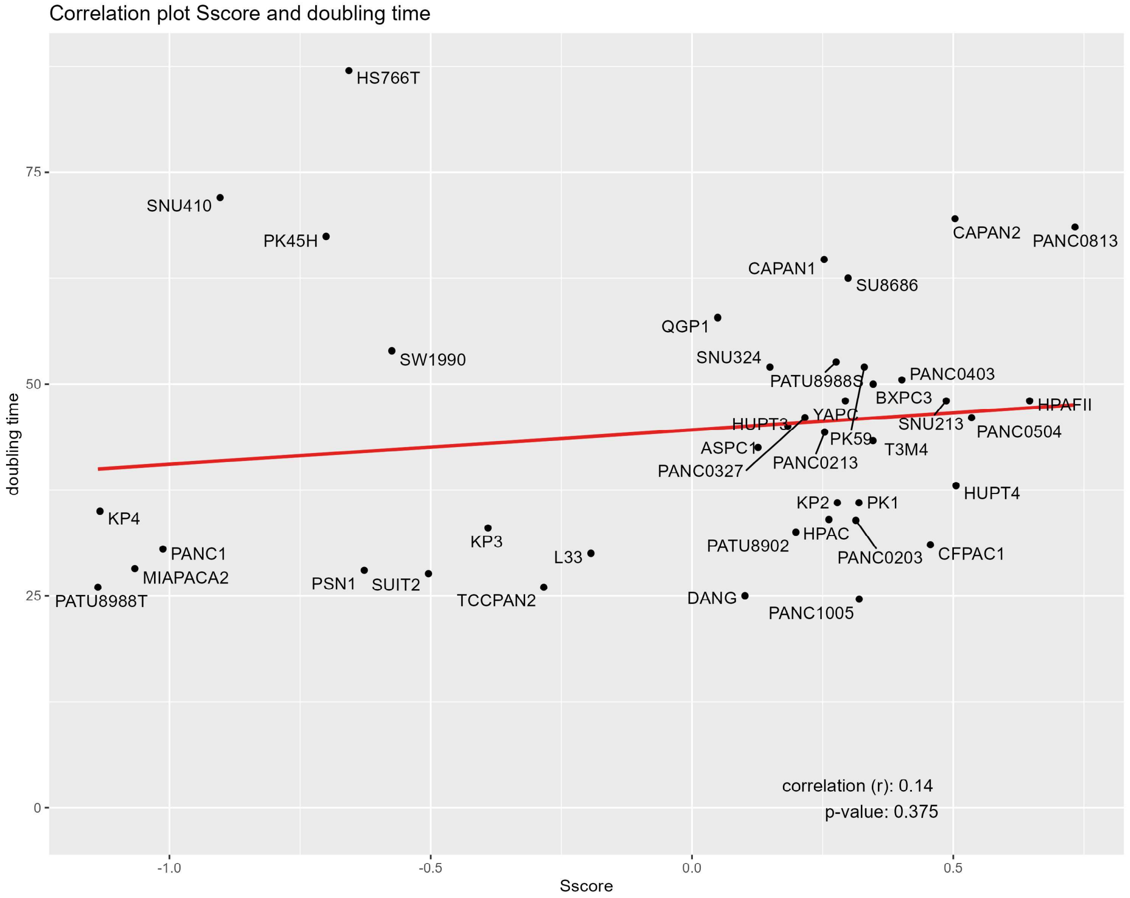

Correlation plot Sscore and doubling time
doubling time
Sscore
0
25
50
75
-1.0
-0.5
0.0
0.5
HS766T
SNU410
PK45H
CAPAN2
PANC0813
CAPAN1
SU8686
QGP1
SW1990
SNU324
PATU8988S
PANC0403
BXPC3
HPAFII
HUPT3
YAPC
SNU213
PANC0504
ASPC1
PK59
T3M4
PANC0327
PANC0213
KP2
PK1
HUPT4
KP4
KP3
HPAC
PATU8902
PANC1
L33
PANC0203
CFPAC1
MIAPACA2
PSN1
SUIT2
PATU8988T
TCCPAN2
DANG
PANC1005
correlation (r): 0.14
p-value: 0.375


Figure S2

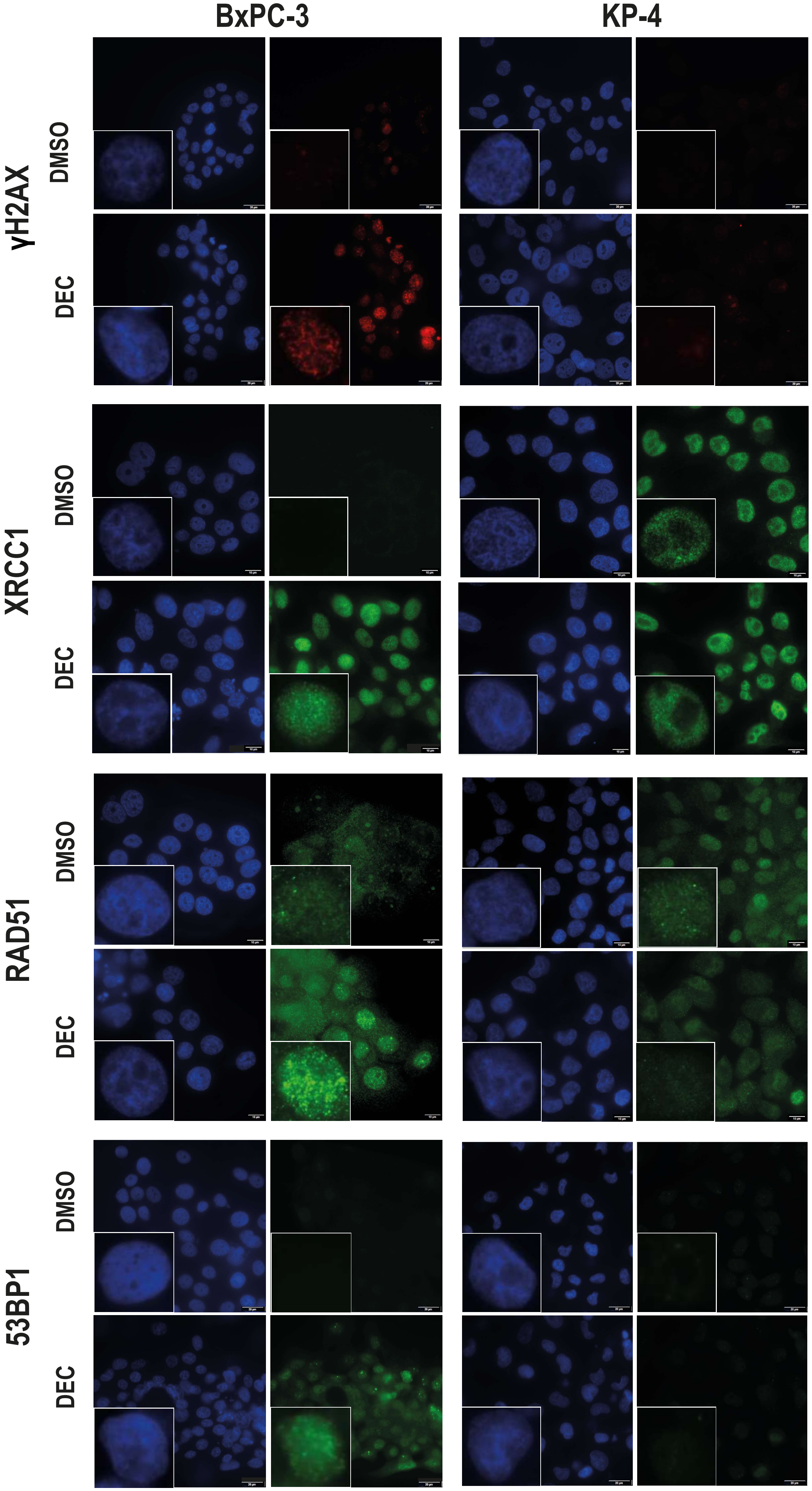
BxPC-3
KP-4
γH2AX
DMSO
DEC
XRCC1
DMSO
DEC
RAD51
DMSO
DEC
53BP1
DMSO
DEC

Figure S3

A

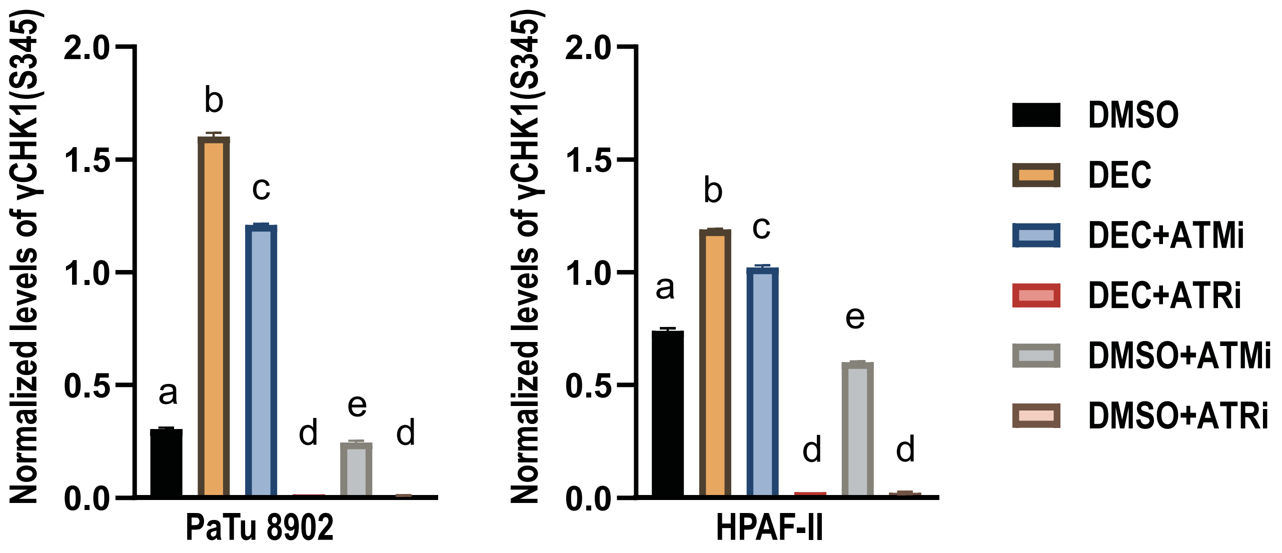
Normalized levels of γCHK1(S345)
2.0
1.5
1.0
0.5
0.0
a
b
c
d
e
d
PaTu 8902
HPAF-II
DMSO
DEC
DEC+ATMi
DEC+ATRi
DMSO+ATMi
DMSO+ATRi

B

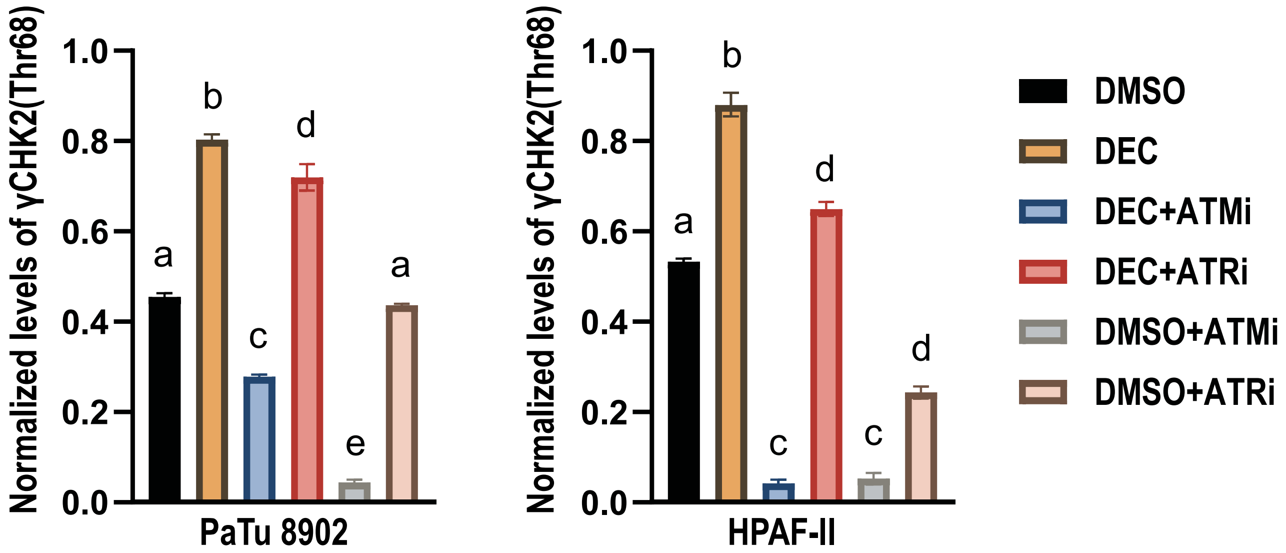
Normalized levels of γCHK2(Thr68)
1.0
0.8
0.6
0.4
0.2
0.0
a
b
c
d
e
a
PaTu 8902
HPAF-II
DMSO
DEC
DEC+ATMi
DEC+ATRi
DMSO+ATMi
DMSO+ATRi

Figure S4

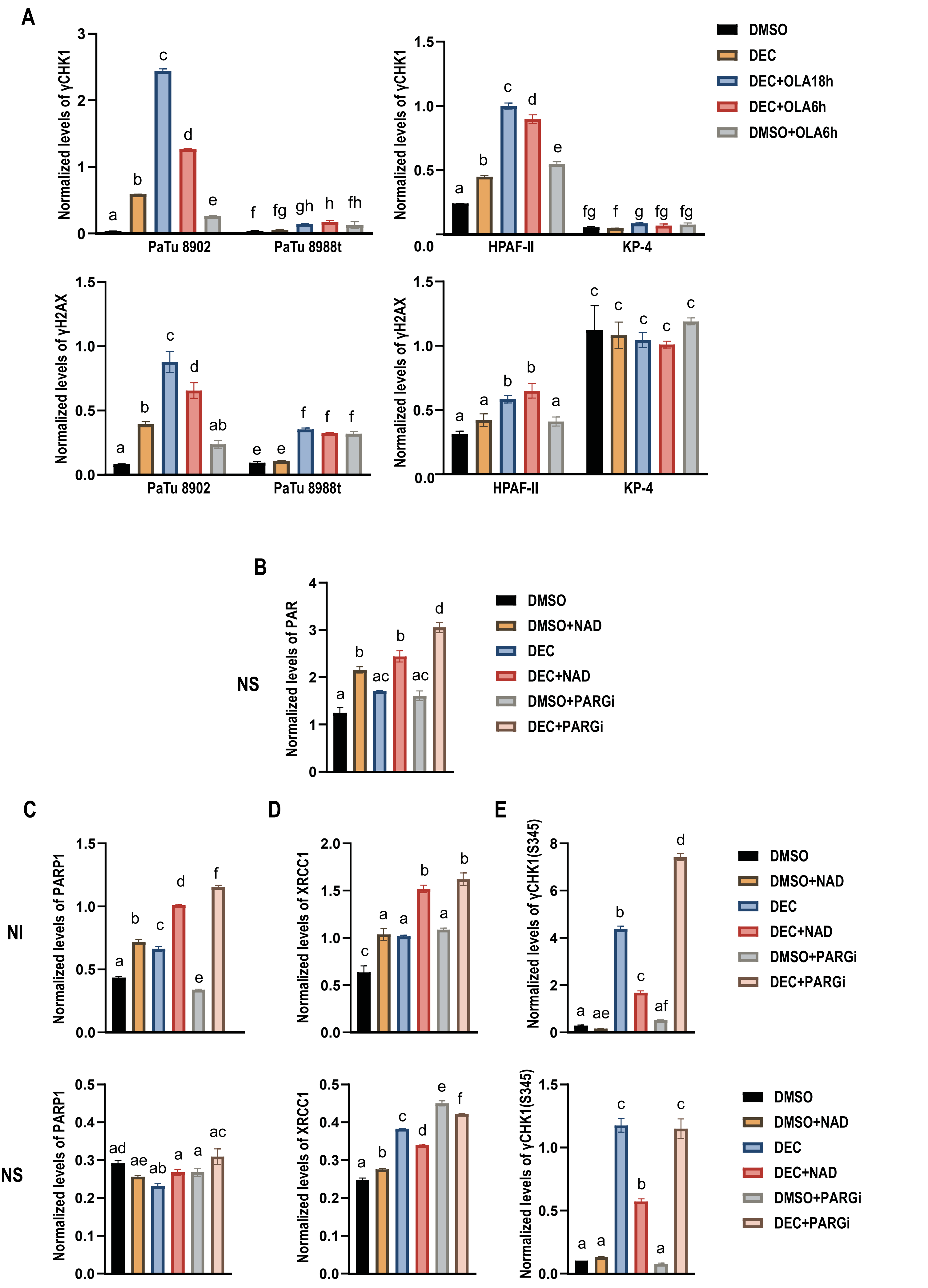
A
Normalized levels of γCHK1
PaTu 8902
PaTu 8988t
HPAF-II
KP-4
Normalized levels of γH2AX
DMSO
DEC
DEC+OLA18h
DEC+OLA6h
DMSO+OLA6h
B
NS
Normalized levels of PAR
DMSO
DMSO+NAD
DEC
DEC+NAD
DMSO+PARGi
DEC+PARGi
C
D
E
NI
NS
Normalized levels of PARP1
Normalized levels of XRCC1
Normalized levels of γCHK1(S345)

Figure S5

| DEC | | |
|---|---|---|
| Cell line | IC50 uM | 95% CI |
| CAPAN-1 | 0.2 | [0.1-0.4] |
| PaTu 8902 | 0.1 | [0.07-0.13] |
| BxPC-3 | 0.15 | [0.1-0.22] |
| HPAF-II | 0.18 | [0.15-0.22] |
| SU8686 | 0.8 | [0.46-1.5] |
| PANC-1 | 1.4 | [0.85-2.4] |
| ASPC1 | 22.9 | [13.8-39] |
| PaTu 8988t | 32.3 | [18.8-61.3] |
| KP-4 | 22.5 | [13.8-44.3] |

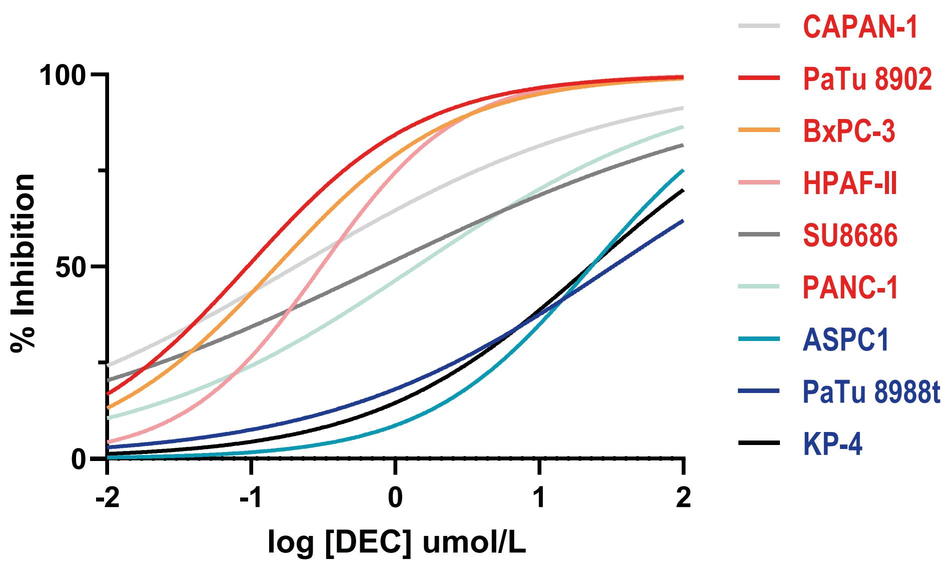


| OLA | | |
|---|---|---|
| Cell line | IC50 uM | 95% CI |
| CAPAN-1 | 0.95 | [0.37-2.2] |
| PANC-1 | 2.6 | [1.7-4] |
| HPAF-II | 5.3 | [4.6-6.2] |
| PaTu 8902 | 5.6 | [4.8-6.6] |
| SU8686 | 7.4 | [5.1-10.9] |
| BxPC-3 | 8 | [5.2-12.6] |
| PaTu 8988t | 9.2 | [7.2-12.1] |
| ASPC1 | 10.1 | [4.6-27] |
| KP-4 | 16.3 | [12.9-20.9] |

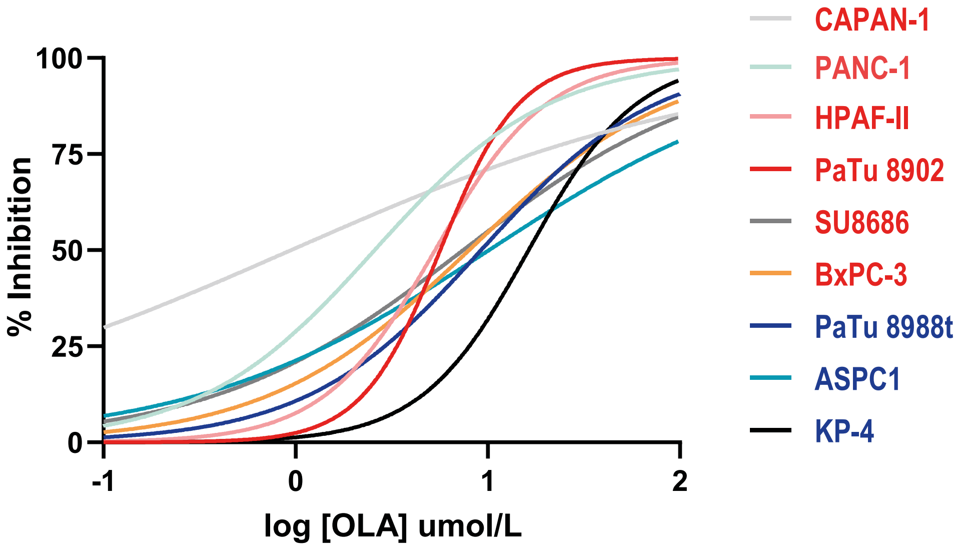


Figure S6

**PaTu 8902**

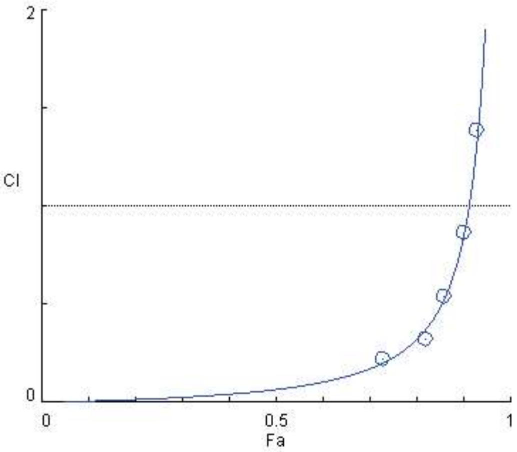


Data for Fa = 0.5

| Drug/Combo | CI value | Dose DEC | Dose OLA |
|---|---|---|---|
| DEC | | 0.11264 | |
| OLA | | | 6.62891 |
| COMBO | 0.06543 | 0.00381 | 0.20953 |

Data for Fa = 0.75

| Drug/Combo | CI value | Dose DEC | Dose OLA |
|---|---|---|---|
| DEC | | 0.29736 | |
| OLA | | | 10.9054 |
| COMBO | 0.22970 | 0.02732 | 1.50286 |

Data for Fa = 0.9

| Drug/Combo | CI value | Dose DEC | Dose OLA |
|---|---|---|---|
| DEC | | 0.78504 | |
| OLA | | | 17.9408 |
| COMBO | 0.85046 | 0.19598 | 10.7791 |

Data for Fa = 0.95

| Drug/Combo | CI value | Dose DEC | Dose OLA |
|---|---|---|---|
| DEC | | 1.51932 | |
| OLA | | | 25.1703 |
| COMBO | 2.12823 | 0.74851 | 41.1678 |

**HPAF-II**

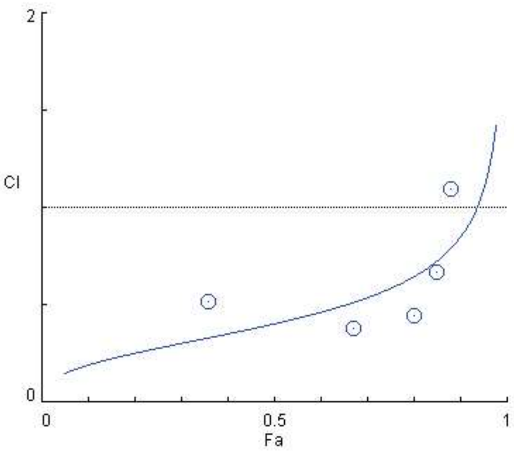


Data for Fa = 0.5

| Drug/Combo | CI value | Dose DEC | Dose OLA |
|---|---|---|---|
| DEC | | 0.36460 | |
| OLA | | | 9.48860 |
| COMBO | 0.40400 | 0.05530 | 2.39433 |

Data for Fa = 0.75

| Drug/Combo | CI value | Dose DEC | Dose OLA |
|---|---|---|---|
| DEC | | 0.89630 | |
| OLA | | | 22.0057 |
| COMBO | 0.58671 | 0.19028 | 8.23920 |

Data for Fa = 0.9

| Drug/Combo | CI value | Dose DEC | Dose OLA |
|---|---|---|---|
| DEC | | 2.20342 | |
| OLA | | | 51.0349 |
| COMBO | 0.85271 | 0.65478 | 28.3522 |

Data for Fa = 0.95

| Drug/Combo | CI value | Dose DEC | Dose OLA |
|---|---|---|---|
| DEC | | 4.06247 | |
| OLA | | | 90.4370 |
| COMBO | 1.10011 | 1.51751 | 65.7082 |

**CAPAN-1**

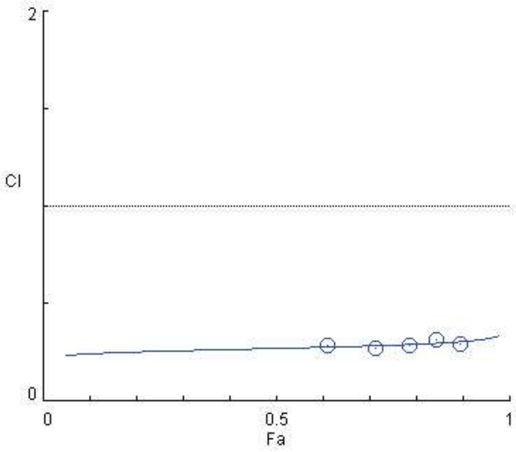


Data for Fa = 0.5

| Drug/Combo | CI value | Dose DEC | Dose OLA |
|---|---|---|---|
| DEC | | 0.08580 | |
| OLA | | | 0.77350 |
| COMBO | 0.27114 | 0.01042 | 0.11578 |

Data for Fa = 0.75

| Drug/Combo | CI value | Dose DEC | Dose OLA |
|---|---|---|---|
| DEC | | 0.46681 | |
| OLA | | | 4.73053 |
| COMBO | 0.28778 | 0.06408 | 0.71198 |

Data for Fa = 0.9

| Drug/Combo | CI value | Dose DEC | Dose OLA |
|---|---|---|---|
| DEC | | 2.53972 | |
| OLA | | | 28.9308 |
| COMBO | 0.30648 | 0.39403 | 4.37816 |

Data for Fa = 0.95

| Drug/Combo | CI value | Dose DEC | Dose OLA |
|---|---|---|---|
| DEC | | 8.03773 | |
| OLA | | | 99.1426 |
| COMBO | 0.32052 | 1.35534 | 15.0593 |

**Figure S7**

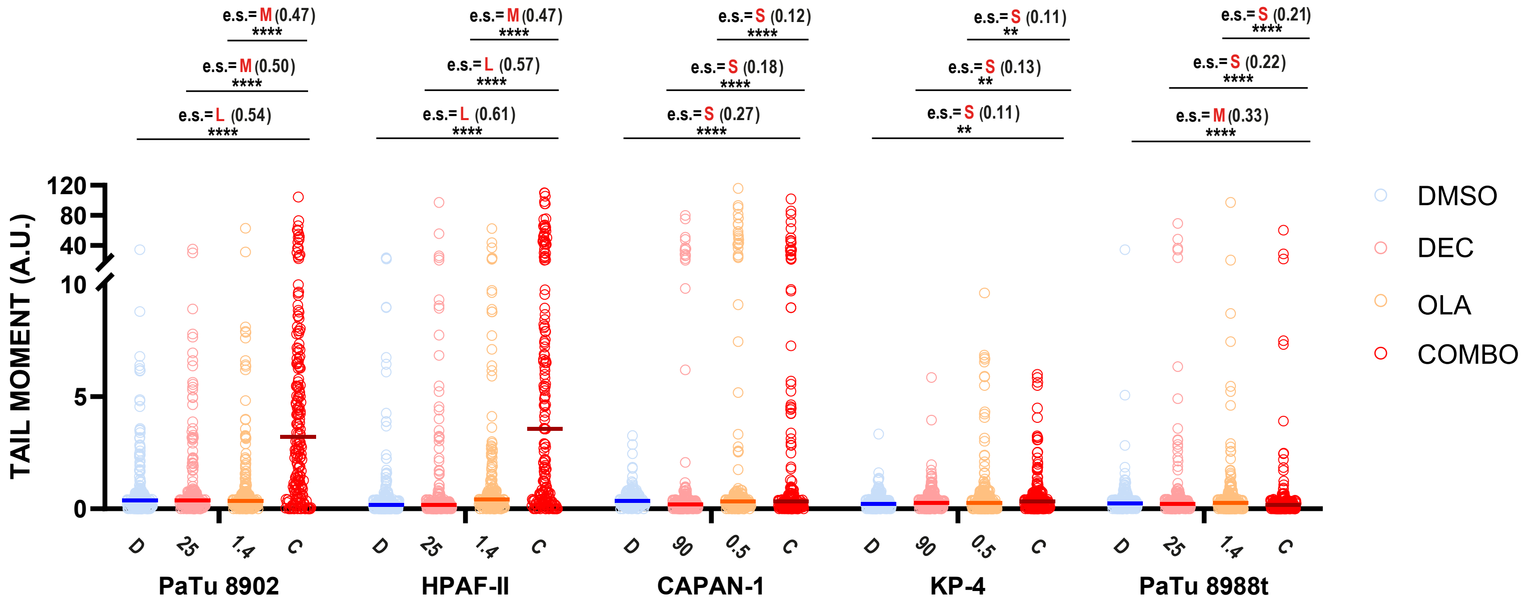

e.s.= M (0.47)
****
e.s.= M (0.50)
****
e.s.= L (0.54)
****
e.s.= M (0.47)
****
e.s.= L (0.57)
****
e.s.= L (0.61)
****
e.s.= S (0.12)
****
e.s.= S (0.18)
****
e.s.= S (0.27)
****
e.s.= S (0.11)
**
e.s.= S (0.13)
**
e.s.= S (0.11)
**
e.s.= S (0.21)
****
e.s.= S (0.22)
****
e.s.= M (0.33)
****
TAIL MOMENT (A.U.)
120
80
40
10
5
0
D
25
1.4
C
PaTu 8902
D
25
1.4
C
HPAF-II
D
90
0.5
C
CAPAN-1
D
90
0.5
C
KP-4
D
25
1.4
C
PaTu 8988t
DMSO
DEC
OLA
COMBO


### Figure S8

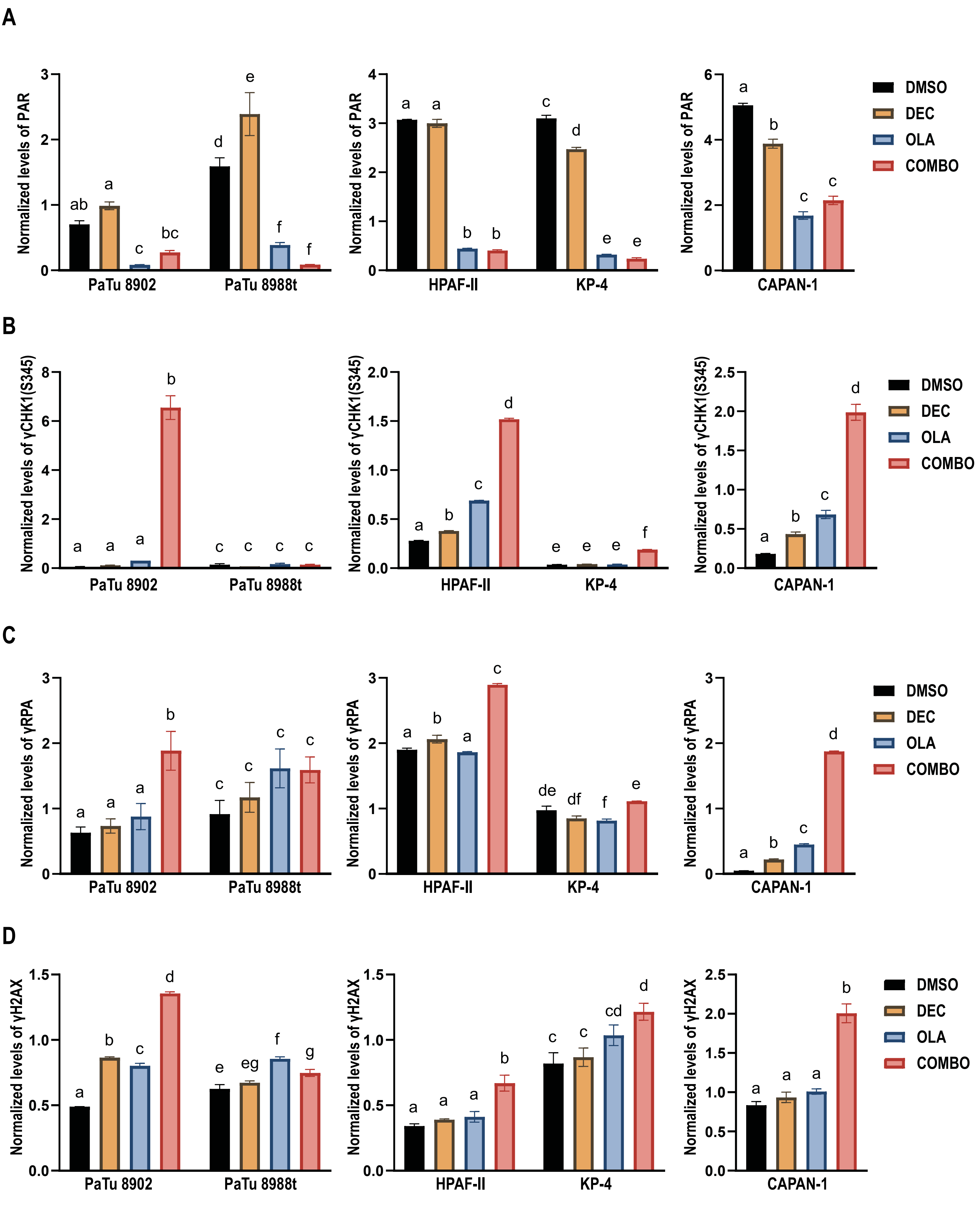
A
Normalized levels of PAR
PaTu 8902
PaTu 8988t
HPAF-II
KP-4
CAPAN-1
DMSO
DEC
OLA
COMBO
B
Normalized levels of γCHK1(S345)
C
Normalized levels of γRPA
D
Normalized levels of γH2AX

Figure S9

A

| | Survival (%) | | |
|---|---|---|---|
| Treatment | DAY 7 | DAY 9 | DAY 28 |
| DEC (0,5 mg/Kg) | 100% | 100% | 80% |
| DEC (0,2 mg/Kg) | 100% | 100% | 100% |
| OLA (30 mg/Kg) | 100% | 100% | 100% |
| OLA (30 mg/Kg) + DEC (0,2 mg/Kg) | 100% | 100% | 100% |

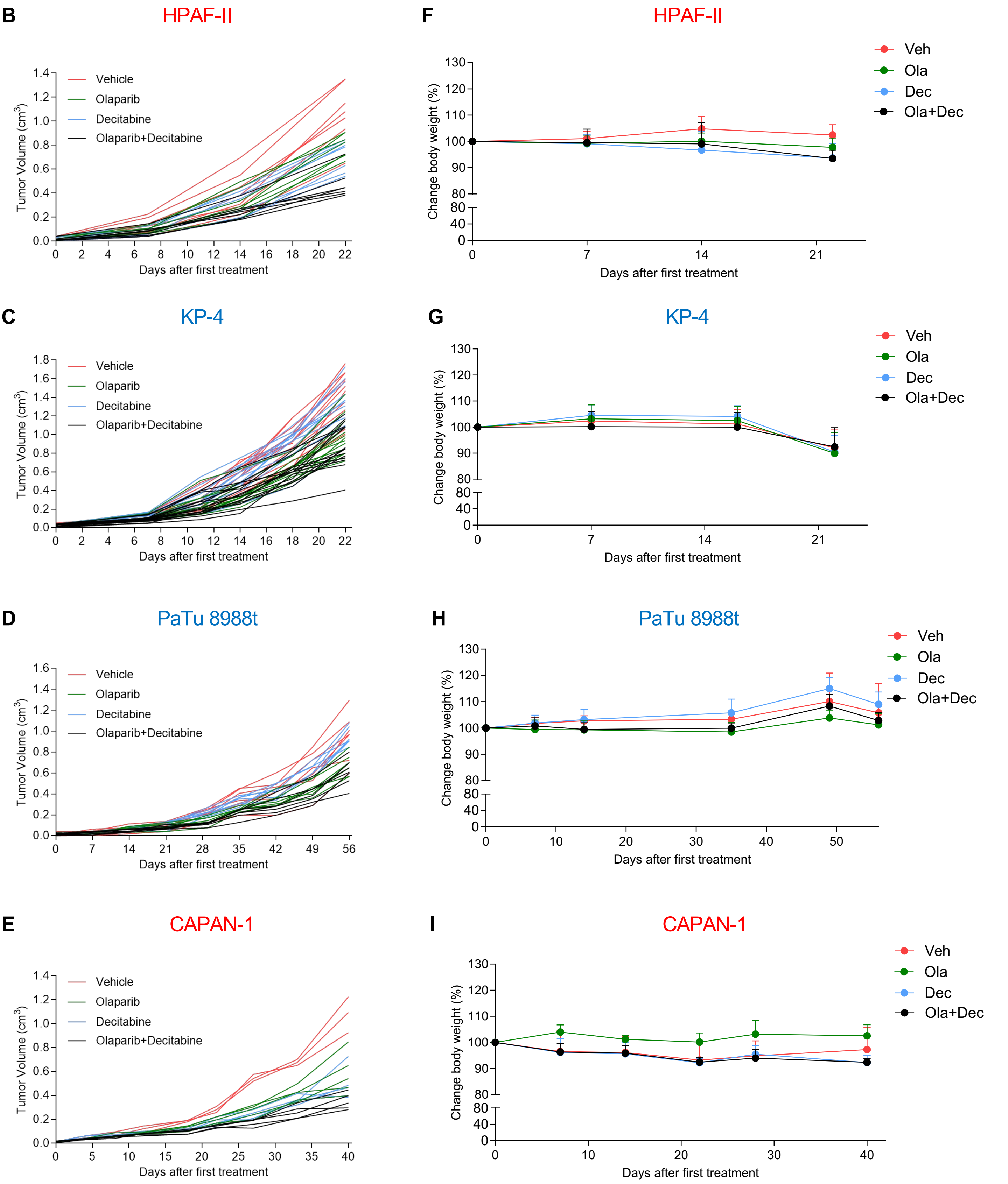


**Figure S10**
